# The transmembrane domain regulates the kinetics of the SARS-CoV-2 spike conformational transition

**DOI:** 10.64898/2026.04.03.716256

**Authors:** Avijeet Kulshrestha, Arkadeep Banerjee, Sahil Lall, Shachi Gosavi

## Abstract

**ABSTRACT:** The homotrimeric SARS-CoV-2 spike glycoprotein comprises two subunits: S1, which recognizes host-receptors through its receptor-binding domains (RBDs), and S2, anchored to the viral membrane through its transmembrane domain (TMD), which facilitates the fusion of the viral envelope with the host cell membrane. Upon host-receptor engagement and proteolytic activation, S1 dissociates and triggers a large conformational transition in S2, involving structural rearrangements in the S2 ectomembrane-domains and the TMD. While studies have focused on the ectomembrane-domain dynamics, the TMD has typically been modeled as being in a trimeric state. Here, we use molecular dynamics simulations of a coarse-grained structure-based model (SBM) with an implicit membrane to investigate the role of TMD dynamics in modulating S2 conformational conversion. We first recapitulate previous results from an all-atom SBM with a trimeric TMD and re-emphasize that the extended pre-hairpin intermediate state of S2, which brings the two membranes into apposition, is a byproduct of the prefusion-to-postfusion transition. Next, by introducing dynamics into the TMD, we find a late fusion intermediate structurally similar to a recent cryo-EM structure. A dynamic TMD also makes the conformational transition faster. Simulations including the S1/S2 complex reveal coupled RBD-TMD dynamics: when all three RBDs are in the closed state, they can stabilize the TMD in a trimeric configuration, whereas the opening of a single RBD can trigger a transition to a dynamic TMD. So, the dynamics and the conformational preferences of the TMD can be tuned by the presence and conformation of S1. There is some evidence that the TMDs of class I viral fusion proteins, such as spike, contribute to viral fusion by modulating membrane properties. Our simulations indicate an expanded role for the function of the TMD, where it can directly regulate the kinetics of S2 conformational transitions.

**SIGNIFICANCE STATEMENT:** The SARS-CoV-2 fusion protein, spike, undergoes a large conformational transition, which facilitates the fusion of the viral and host-cell membranes and the delivery of the viral genome into the host cell. Despite extensive studies of the spike conformational conversion, its transmembrane domain (TMD) has largely been viewed as a viral membrane anchor. Using coarse-grained structure-based model simulations, we show that TMD dynamics can modulate the timing of the spike-S2 (fusion subunit) prefusion-to-postfusion conformational conversion. The presence of spike-S1 (receptor-recognition subunit) can suppress TMD dynamics, potentially reducing the rate of spike conformational conversion and viral fusion. Thus, the spike TMD regulates the kinetics of spike-mediated membrane fusion, and TMD-targeting strategies can be an additional avenue for antiviral intervention.

## INTRODUCTION

The homotrimeric spike (S) glycoprotein mediates the fusion of the viral membrane of Severe Acute Respiratory Syndrome Coronavirus 2 (SARS-CoV-2), an enveloped RNA virus responsible for the COVID-19 pandemic, with the host cell membrane (**Figure 1**). The fusion of the two membranes facilitates the delivery of the viral genome into host cells (1). A furin cleavage motif within the S protein sequence, absent in closely related coronaviruses (2, 3), is cleaved during protein biosynthesis to produce the S1 (residues 1-685) and S2 (residues 686-1273) subunits, which remain noncovalently associated (**Figure 1**) (4). The S1 subunit mediates host cell recognition by interacting with the host angiotensin-converting enzyme 2 (ACE2) receptor via its flexible receptor-binding domain (S1-RBD), which fluctuates between closed and open conformations, with the latter conformation being prone to receptor engagement (5). The S2 subunit, comprising the fusion peptides (FP), heptad-repeat regions HR1 and HR2, transmembrane domain (TMD), and cytoplasmic tail (CT), plays a central role in membrane fusion (**Figure 1**). The binding of the S1-RBD to ACE2 is accompanied by a proteolytic cleavage at the S2’ site (residue S816) within the S2 subunit, which is mediated either by the host cell membrane anchored serine protease TMPRSS2 or by cathepsin L within endo-lysosomes (1). Together, these events enable the shedding of the S1 subunit and expose the fusion peptide (FP), triggering the transition of the cleaved S2 subunit from the prefusion to the postfusion conformation (6). This S2 conformational transition and the intermediates populated during this transition facilitate the fusion of the virus and host membranes. Several experimental studies have attempted to characterize these spike intermediates by arresting them through different approaches (7–12). However, most of these studies have focused on the large ectomembrane domains of the spike.

**Figure 1:**
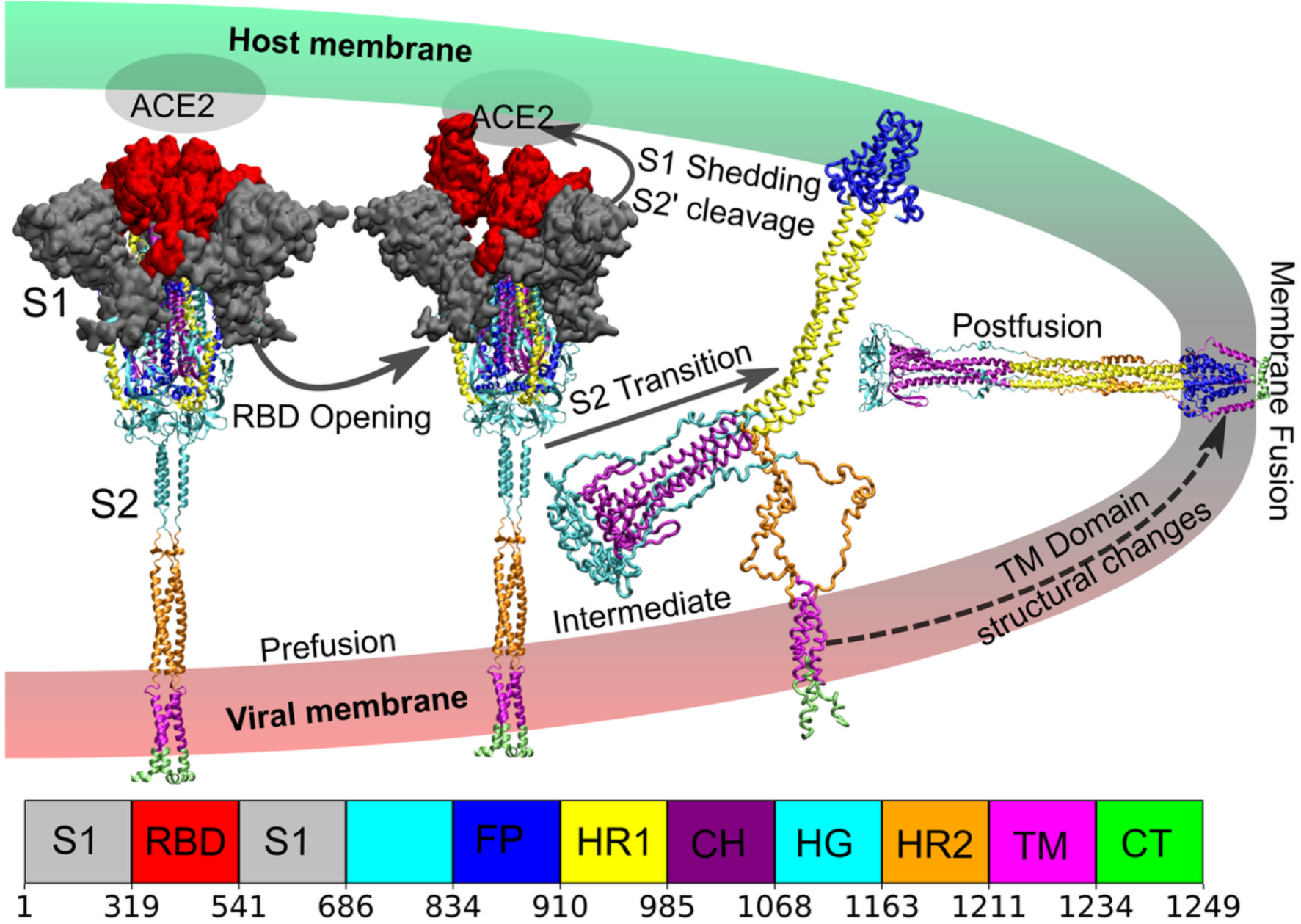
Spike (S) protein conformational transition. The structures of spike during the conformational transition (above) are colored according to the domain architecture with residue numbering (below). The furin cleavage at the S1/S2 boundary (R685) produces the S1 (shown in surface representation) and S2 (shown in cartoon representation) subunits. During the transition of the S protein from the prefusion (1^st^ structure) to the postfusion state (last structure, drawn horizontally), the opening of the receptor-binding domain (RBD) of the S1 subunit leads to interaction with the human host ACE2 receptor (2^nd^ structure), promoting S1 shedding, S2’ cleavage at residue S816, and triggering the S2 conformational change. S2 subsequently adopts an extended prehairpin intermediate (3^rd^ structure; snapshot extracted from the simulations, where the S2 (R686)-S2’ (S834) segment was not modeled in the simulations and is not shown). The fusion peptide docks with the host membrane, and subsequent conversion of S2 into the postfusion state brings the viral and host membranes into close proximity, ultimately resulting in membrane fusion. The transmembrane domain (TMD; pink and marked as TM in the domain architecture) consists of three single pass transmembrane helices. The structural changes in the TMD are shown with a dashed arrow within the viral membrane. A detailed version of the S2 domain description is given in **Figure S2A**.

As stated earlier, the S2 subunit contains the TMD, which is comprised of three single-pass transmembrane helices, each contributed by one of the protomers of the homotrimeric spike. Owing to its membrane localization and small size, the TMD has been difficult to study experimentally. Therefore, the TMD is generally assumed to form a three-helix bundle and function as an anchor that tethers the large spike protein to the viral membrane without affecting the S2 conformational transition and viral fusion. Although mutations in the conserved Trp-rich regions of the TMDs of SARS-CoV and SARS-CoV-2 do not prevent spike expression or RBD binding, they have been shown to affect trimer stability and infectivity (13–15), suggesting a functional role for the TMD beyond passive anchoring. Further, an examination of the SARS-CoV-2 spike postfusion structure (16) shows that the TMD likely undergoes a structural rearrangement and must open from its potentially trimeric prefusion conformation to wrap around the fusion peptides (FPs) in the postfusion state (**Figure 1**). Additionally, molecular dynamics simulations have shown that the helices that comprise the TMD are highly flexible and undergo conformation-dependent trimerization in POPC membranes, which is reduced in the presence of cholesterol (17). Thus, there are clear indications that the TMD is dynamic and functional. Here, we investigate the role of TMD dynamics in modulating the spike-S2 conformational transition using molecular dynamics (MD) simulations of structure-based models (SBMs), coarse-grained to a single bead per residue located at the Cα position.

Structure-based models (SBMs) encode the folded or native structure of the protein in their potential energy functions and have been successfully used to study large-scale protein folding and conformational transitions, reproducing experimental kinetics and providing mechanistic insights (18–22). SBMs are particularly valuable when atomistic molecular dynamics simulations are computationally challenging due to system size and timescale (23). Recently, all-atom SBMs were used to study the conformational conversion between early SARS-CoV-2 spike pre- and post-fusion structures (8, 11). These simulations showed that glycan post-translational modifications could sterically hinder the spike conformational change, providing a critical pause for fusion peptides to engage the host membrane (11). These all-atom SBM simulations were initiated from the pre-fusion structure with the potential energy function encoding only the post-fusion structure, thus promoting conformational conversion. However, the TMD was constrained to be a helical bundle throughout the simulations, making it difficult to study its dynamics. Additionally, after the all-atom SBM simulations were performed, a near complete post-fusion structure of the SARS-CoV-2 spike was experimentally resolved, which includes the membrane-embedded TMD-fusion peptide complex (16). Here, we perform simulations using an SBM coarse-grained to a single Cα-bead per residue. These simulations were performed in a manner analogous to the previous all-atom SBM simulations of spike (11), i.e., they are initiated from (partially modeled) prefusion structure. The recent experimentally resolved post-fusion structure (16) is encoded in the potential energy function. Our simulations also have an implicit membrane representation; however, we do not constrain the TMD to be a trimeric helical bundle. We first show that the Cα-SBM simulations show a mechanism of conformational conversion similar to that of the all-atom SBMs, and as with those simulations, we find that the extended host-membrane interacting pre-hairpin intermediate of S2 (**Figure 1**) is a by-product of the conformational conversion and does not need to be encoded into the potential energy function. Additionally, we find the population of a late fusion intermediate, which is structurally similar to a recently resolved cryo-EM structure (24). We then construct a variant Cα-SBM that encodes the postfusion state (as before) but with a trimeric TMD stabilized using Lennard-Jones-like interactions, which can break and form during the simulation. Simulations of this Cα-SBM variant slow down the conformational conversion from the prefusion to the postfusion state, and there is a correlation between TMD dynamics (how early the TMD interactions break) and the kinetics of the conformational transition. Thus, TMD dynamics are important for the timing of the S2 conformational conversion.

Several studies have suggested that the S1-RBD movement from a closed state to an open state, potentially upon host receptor engagement, may result in S1 shedding and trigger the S2 conformational transition (8, 25, 26). Since S1 is far removed from the TMD, it is unclear whether the S1-RBD opening in the spike pre-fusion state, leading to a loss of S1/S2 interactions, could affect TMD dynamics. To answer this, we simulated S2 with the stabilized trimeric TMD in the presence of S1 and examined TMD dynamics. We find that the presence of S1 increases the population of the trimeric TMD, with the S1-RBD in the closed state having a higher trimeric TMD population than S1 with one open RBD. Together, the simulations indicate that TMD dynamics can play an important role in the kinetics of the spike conformational change, and that these TMD dynamics are themselves modulated by the conformational state of the RBD.

## RESULTS

### Cα-SBM captures the spike transition

We simulated 100 independent trajectories of the prefusion-to-postfusion transition initiated from distinct prefusion structures (see Methods). Only the postfusion structure was encoded in the Cα-SBM potential energy function, and no information from the prefusion structure was included. An implicit membrane region was defined using a reverse flat-bottom potential (see Methods), which is applied to the Z-coordinate of membrane abutting regions (HG, HR2, and CT domain residues; **Figure 1**), preventing them from entering the membrane and indirectly keeping the TM domain (TMD) within the membrane region.

To analyze conformational conversion during the simulations, we first compute the root-mean-square deviation (RMSD) of each simulation frame with respect to the postfusion structure (**Figure S3B**). The RMSD initially increases to ∼10.7−12.8 nm for all trajectories due to the population of the extended pre-hairpin conformation (docking of this conformation with the host membrane brings the two membranes into apposition), after which the RMSD decreases and approaches zero within ∼2×10^7^−7.7×10^7^ steps. To further characterize the conformational changes and for ease of comparison with previous all-atom SBM simulations (11), we defined three structural parameter derived from those used in previous simulations (**Figure 2A and 2C**): FP_Z_, the Z-coordinate of the distance between the center-of-mass of fusion peptides and the TM domain; Z_head_, the Z-coordinate of the center of mass of the head (HG) relative to the TM domain; and r_head_, the lateral distance between the head and the TM domain. The FP_Z_ values (**Figure 2B**) show that the fusion peptides first extend relative to the TM domain up to ∼30 nm, corresponding to the first intermediate state (I1; **Figure 2A**), which is similar to the extended pre-hair intermediate reported in prior studies (7–12). Notably, this intermediate arises naturally in the SBM simulations (also see (11)) and is not explicitly encoded in the potential energy function. This conformation bridges the viral and host cell membranes (**Figure 1**) because the fusion peptides reach out to interact with the host membrane while the TM helices remain anchored in the viral membrane. Following this extension, FP_Z_ abruptly decreases as HR2 zips onto HR1, producing a second intermediate state, I2, that closely resembles the postfusion conformation. Such cooperative zipping has been shown to be a common mechanistic feature across several class-I viral fusion proteins (27). A recently resolved late-fusion cryo-EM intermediate structure closely resembles I2 (**Figure 2C**) (24).

**Figure 2:**
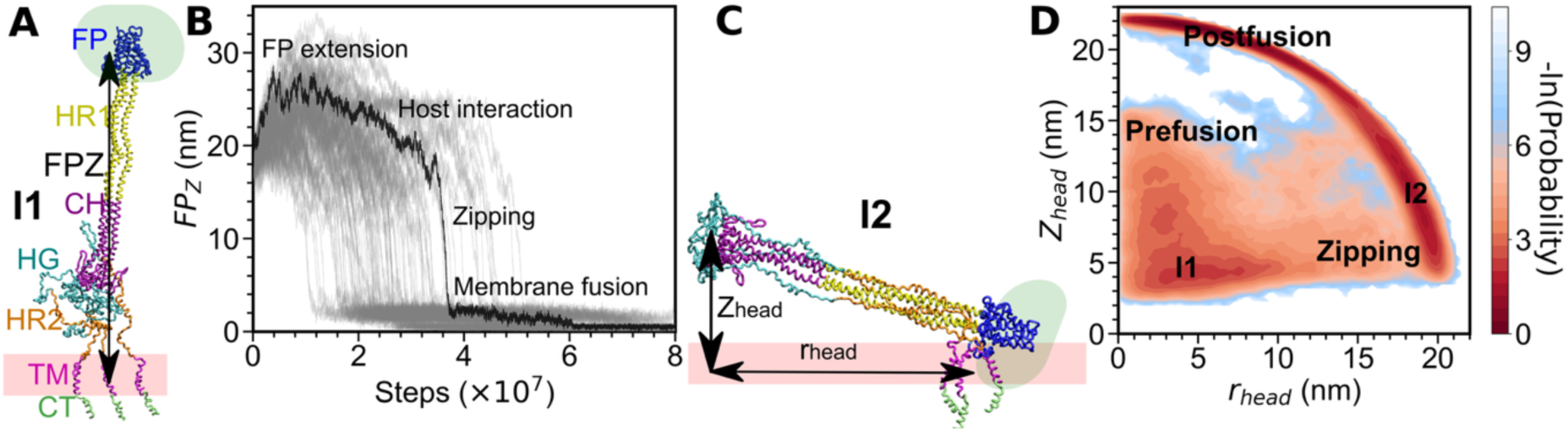
Spike transition mechanism from the unrestrained TMD model encoding only the postfusion structure. (A) Structure of the spike in the I1 intermediate. FP_Z_, the difference between the Z-coordinates of the centers-of-mass of the transmembrane domain (marked TM) and the fusion peptide (FP), is marked. FP_Z_ can be used to quantify the FP extension and other features of the FP dynamics. (B) FP_Z_ from individual simulations (100 replicates), each plotted with a gray line as a function of time. One simulation trajectory is highlighted in black and specific structural events are labelled. (C) Structure of the spike in the I2 intermediate. Two structural metrics which quantify the motion of the head region (HG) are marked. Z_head_ is the difference between the Z-coordinates of, and r_head_ is the radial (along the membrane) distance between, the centers of mass of the TMD and HG regions. (D) Two-dimensional free-energy landscape with Z_head_ and r_head_ as coordinates. The transition pathway between the prefusion structure through I1, zipping and I2 to the postfusion structure can be seen. The two representative intermediate structures in (A):I1 and (C):I2 are colored as in Figure 1.

To further examine the transition mechanism, we plotted a 2D “free-energy” surface using r_head_ and Z_head_ (**Figure 2D**; since these are transition simulations, -ln(probability) is only analogous to the free-energy of equilibrium simulations; see Methods), which shows the Z_head_ first approaches ∼5 nm with minimal change in r_head_, implying a head (HG) movement towards the TM domain to reach the first intermediate (I1; **Figure 2A**) state. This downward motion facilitates fusion-peptide exposure and extension toward the host membrane. The system further progresses to the second intermediate state (I2; **Figure 2C**), where Z_head_ increases slightly and r_head_ increases to ∼20 nm, indicating substantial lateral movement of the head that brings the host and viral membranes closer together. Finally, the TM domain accommodates the fusion peptide as the protein changes both Z_head_ and r_head_ to reach the postfusion state. The overall mechanism is consistent with the previous all-atom SBM results (11). However, the all-atom SBM simulations did not have the I2-to-postfusion transition because the experimental postfusion structure had not been characterized at that time and the SBM lacked TMD-FP interactions.

The two intermediate states, I1 and I2, are strikingly similar to recently resolved early (12) and late (24) fusion intermediates. Overall, the Cα-SBM simulations of spike-S2 can provide insight into the mechanism of the pre-fusion to post-fusion conformational conversion and also recapitulate some of the experimentally observed intermediates. We next set out to understand the effect of TMD dynamics on this conformational transition.

### TMD dynamics regulate the spike-S2 transition kinetics

The prefusion atomistic structure of the TMD in the context of the full-length spike protein has not been structurally resolved either due to its small size, membrane embedding, or dynamics (17). However, the structure of the isolated TMD (with several mutations), characterized using NMR (28), is a symmetric trimer. Given this, we simulated two extreme cases: (i) <u>Unrestrained TMD</u>: Only the postfusion structure is encoded in the Cα-SBM (see previous section). Since very few stabilizing inter-helical contacts are present between the TMD helices in the postfusion structure, the TMD helices can move freely within the membrane, constrained only by the ectomembrane regions of the spike-S2. (ii) <u>Trimeric TMD</u>: The Cα-SBM is modified to include stabilizing inter-helical TMD contacts (111 in number) extracted from its NMR structure (28). These contacts are modeled using a Lennard-Jones-like potential and can break and form during the simulations. This model is similar to the model used in the previous all-atom SBM simulations of spike-S2, except that those TMD contacts were modeled using a harmonic potential (11) and could not break. In this section, we describe the spike-S2 simulations with the trimeric TMD and compare them with the unrestrained TMD spike-S2 simulations from the previous section.

The additional NMR-derived contacts stabilize a trimeric TMD population, which needs to unfold in order to form the postfusion TMD-FP interactions (**Figure 1**; TMD structural changes). Overstabilized TMD contacts can suppress their unfolding, making it difficult to form the TM-FP interactions and transition to the postfusion state. To prevent this, we studied the spike-S2 conformational transition at different interaction strengths (1.0, 0.9, 0.8 times the interaction strength of other contacts) of the NMR derived TMD contacts and quantified both the fraction of inter-helical TMD contacts and the fraction of the TM-FP (post-fusion) contacts. The 2D free-energy surfaces (**Figure S4**) indicate that at interaction strengths above 0.8, the prefusion contacts overstabilize the TM domain, preventing many trajectories from reaching the postfusion state. Therefore, we chose to study the spike-S2 conformational transition with NMR-derived TMD contacts at 80% of the strength of all other contacts (contacts derived from the postfusion structure).

To compare with the unrestrained TMD simulations **(Figure 2**; contacts derived only from the postfusion state), we simulated 100 independent transitions of the model with the trimeric TMD and computed the RMSD relative to the postfusion state. A comparison of the RMSD analysis for the two models (**Figure S5A**) and of their mean RMSD (**Figure S5B**) shows that a trimeric TMD delays the conformational transition. We further computed FP_Z_ (the Z-coordinate of the distance between the center-of-mass of fusion peptides and TM domain), Z_head_ (the Z-coordinate of the center of mass of the head (HG) relative to the TM domain), and r_head_ (the lateral distance between the head and the TM domain) for the simulations with the trimeric TMD model to compare these metrics with the unrestrained TMD simulations (**Figures 2A and 2C**). A comparison of the 2D free-energy surfaces using r_head_ and Z_head_ for the unrestrained TMD (**Figure 2D**) and trimeric TMD (**Figure S5C**) shows that the mechanism of conformational conversion is similar in the two models. However, the plot of FP_Z_ vs. time (**Figure 3A**) shows that the fusion peptides extend farther (up to a maximum of 36.2 nm across trajectories) for the trimeric TMD than for the dynamic TMD (up to a maximum of 34.8 nm). The FP_Z_ values for all the trajectories of each model were further binned into windows of 10^7^ steps each, which show an approximately bimodal distribution (**Figure 3B)** with one population encompassing the pre-fusion and extended pre-hairpin conformations, while the other population contains all near-post-fusion structures. The extended conformation can be seen to persist longer in the trimeric TMD model, whereas FP_z_ decreases more rapidly when in the unrestrained TMD model. We next quantified the number of steps required for the FP_z_ to reach the postfusion state (i.e., FP_z_ ≤ 0.67 nm) and compared the distributions of the two models (**Figure 3C)**. The trimeric TMD model takes longer to convert to the postfusion state, with the distribution maximum shifted to the right by ∼1×10^7^ steps relative to the unrestrained TMD model.

**Figure 3:**
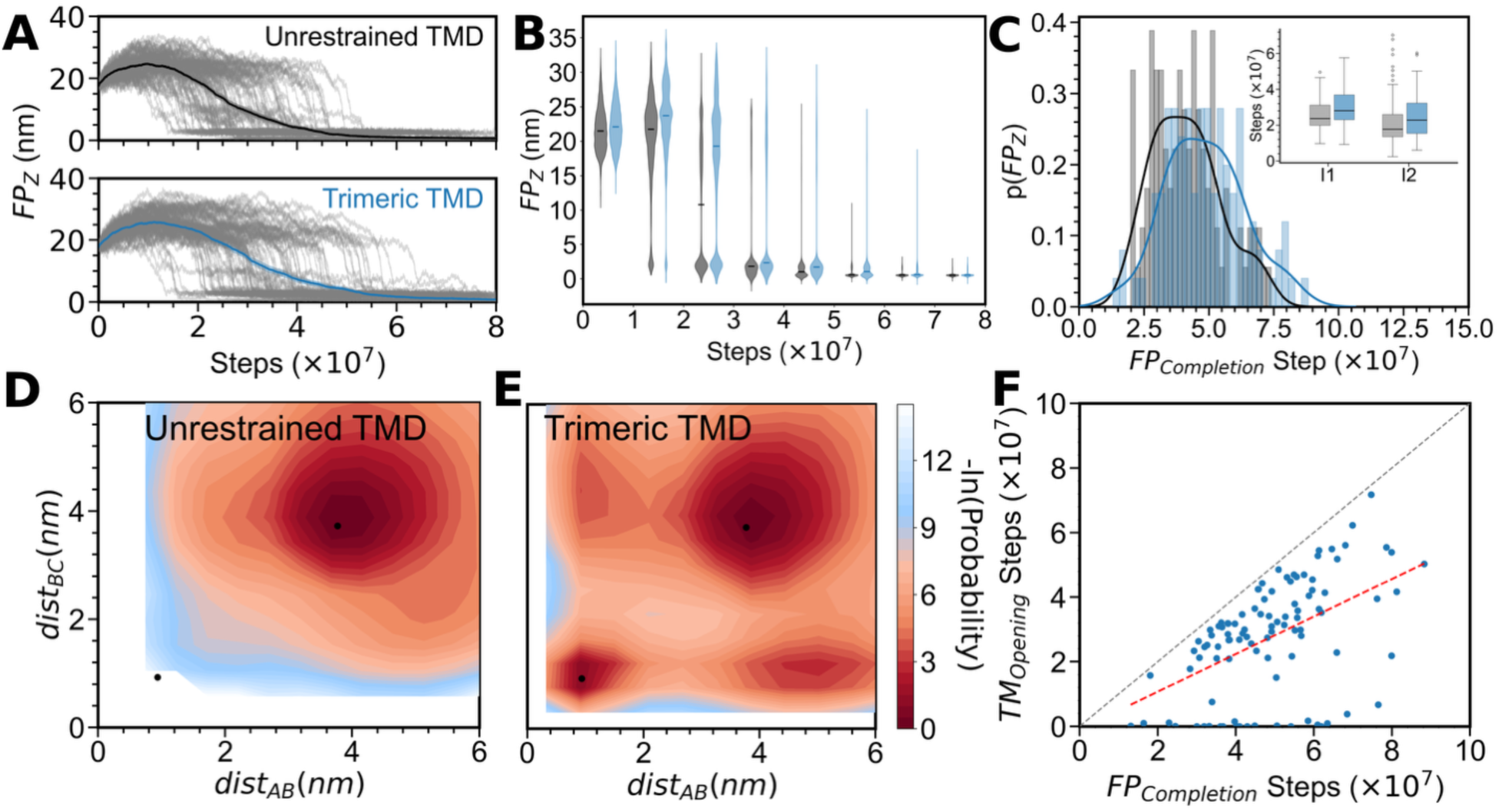
A comparison of the Unrestrained TMD and Trimeric TMD models. (A) Comparison of FP_Z_ (Figure 2A) from the unrestrained (upper panel; data reproduced from Figure 2B) and trimeric TMD (lower panel) models. Individual simulation data (100 replicates; gray lines) plotted as a function of time. The mean is shown. Black/grey is used to depict the unrestrained TMD and blue to depict the trimeric TMD models in (B) and (C). (B) FP_Z_ values from all simulations are pooled for every 10^7^ steps, and a violin plot is shown for each pool. A comparison of the plots from the two models shows that the FP in the trimeric TMD model extends to higher Z values and remains extended for a longer duration than in the unrestrained TMD model. (C) Probability density distribution of the number of steps required for FP_Z_ to reach the postfusion state (FP_completion_; see Results). The inset shows box plots with the median and interquartile ranges for the steps spent sampling the intermediates I1 (Figure 2A) and I2 (Figure 2C) in each trajectory. (D-E) Two-dimensional free-energy surfaces (2D-FES)s with the center-of-mass distance between the TM helix of chain A and chain B and the center-of-mass distance between the TM helix of chain B and chain C as the two coordinates. The color bar is from maroon (high populations; deeper basin) through pale red to blue (low population). Thick dots mark the prefusion structure (near the lower right corner) and the postfusion structure (nearer the upper left corner) distances. (D) Unrestrained TMD model has a single basin close to the postfusion structure. (E) Trimeric TMD model has four basins, two with a single helix dissociated from the trimer. (F) For the trimeric TMD model, the number of steps required for TM domain opening (TM_opening_), defined as the change in interhelical distance from the prefusion to the postfusion state, is plotted vs FP_completion_ for each replicate. The gray dashed line represents the diagonal, and the red dashed line indicates a linear fit to the data (Y = 0.61X - 0.05, R^2^ = 0.32).

The delay in transition kinetics of the trimeric TMD model likely changes the residence times of the two intermediate states, I1 and I2. To quantify this, we defined the following criteria: the protein is considered to populate I1 if Z_head_<15 nm and r_head_<10 nm, and it is considered to populate I2 if Z_head_<15 nm and r_head_>10 nm. The changes in Z_head_ and r_head_, and the associated transitions to I1 and I2, are illustrated for a representative trajectory in **Figure S6A**. Inset box plots in **Figure 3C** compare the timesteps spent in intermediates I1 and I2 (corresponding distributions are shown in **Figures S6B and S6C)** and show that the trimeric TMD model samples both I1 and I2 longer than the unrestrained TMD model (Mann-Whitney p-value 0.0004 and 0.04 for I1 and I2, respectively).

We next compare the TMD behavior in the two models by computing interhelical distances between the helices of the protomers of the TMD (residues 1211 to 1234). The unrestrained TMD model samples large average interhelical distances (>10 nm; **Figure S7B)** due to the absence of energetic stabilization of the trimer. As expected, the trimeric TMD model, with its additional NMR-derived contacts, fluctuates less, with interhelical distances mostly remaining below 5 nm. To further quantify differences in the TMD dynamics, we computed 2D free energy plots of the center-of-mass distances between TMD helices of chain A and chain B (dist_AB_), and between helices of chain B and chain C (dist_BC_). As expected, in the unrestrained TMD model, the TMD has a single population basin whose minimum is near the postfusion state (**Figure 3D**). In contrast, the model with the trimeric TMD first samples the prefusion state before transitioning to the postfusion state (**Figure 3E**). Interestingly, there is a second basin populated when one helix dissociates from the trimer while the other two helices continue to interact. So, helix dissociation is not an all-or-none transition, corroborating previous isolated TMD Martini coarse-grained simulation results (17).

Finally, we test the correlation between the TMD dynamics and conformational conversion of spike protein in the model with the trimeric TMD. We plot the number of steps required for the TMD to interact with the FP (FP_Z_ ≤ 0.67 nm; a proxy for the completion of conformational conversion; **Figure 1**) versus the number of steps required for a trimeric TMD structure to dissociate. TMD dissociation is denoted as TMD_opening_ and is defined as the point at which the average distance between the centers of mass of the three TMD helices (interhelical distances) exceeds the prefusion state interhelical distance (0.93 nm) and remains above this threshold (see **Figure S7A** and method section for the definition). The data suggest that the TMD first opens, followed by the FP transitioning to the postfusion state, as indicated by the presence of data in the lower triangular region (**Figure 3F**). Furthermore, except for trajectories in which the TMD opens rapidly or is initiated from the open conformation, the data show a linear correlation, indicating that the longer it takes for the TMD to open, the longer the conformational conversion takes.

### S1-RBD opening can modulate S2-TMD populations

Both models used in the previous sections assume that the S2 subunit behaves as a “loaded spring” which is released upon S2’ cleavage and S1 dissociation. Upon release, this “stressed” or loaded S2 converts to the postfusion resting state. The presence of the S1 subunit could lock the spring in the loaded form, and its release may depend on the S1 conformation. The S1 subunit can adopt multiple conformations, with all RBDs closed or one or more RBDs open. Host cell ACE2 receptors have been shown to cross-link spike proteins, which can facilitate the opening of multiple RBDs (8). Cryo-EM indicates that RBDs exist in a dynamic equilibrium between closed and open conformations, and binding to the host ACE2 receptor leads to the loss of certain S1-S2 contacts that stabilize one or more open RBDs (25). Overall, RBD opening may, in itself, lead to a partial release of the “loaded” S2 through the loss of S1-S2 contacts.

Therefore, both the presence and conformation of S1 (open or closed RBD) and TMD dynamics (as seen in the previous sections) may modulate the kinetics of the S2 conformational transition. Here, we test if S1 can modulate TMD dynamics by stabilizing the prefusion trimeric conformation and increasing its population. This, in turn, could amplify the population of the prefusion-S2 state (similar to locking S2 in its prefusion structure). We constructed Cα-SBMs of the full spike protein, excluding residues 686 to 833 (sequence between the S2 and S2’ cleavage sites), i.e., assuming that both cleavages had occurred (**Figure 1**). The full spike Cα-SBMs were built as follows: The previous trimeric TMD model (with the postfusion S2 structure) was used for S2 (**Figure 4A**). The S1 region (pre-S2 cleavage site sequence) had either (i) all three RBDs in the closed conformation (this full spike Cα-SBM will henceforth be referred to as S1-closed RBD) or (ii) one RBD in the open conformation (spike Cα-SBM henceforth referred to as S1-open RBD). The S1 region (both S1-closed RBD and S1-open RBD) was modeled using the respective pre-fusion structures (**Figure 4A)**. Inter-(S1-S2) contacts were also derived from the respective pre-fusion structures. The S1-closed RBD Cα-SBM had 355 contacts between S1 and S2, while the S1-open RBD Cα-SBM had 291 contacts between S1 and S2, corresponding to an ∼18% loss of S1-S2 contacts upon RBD opening.

**Figure 4:**
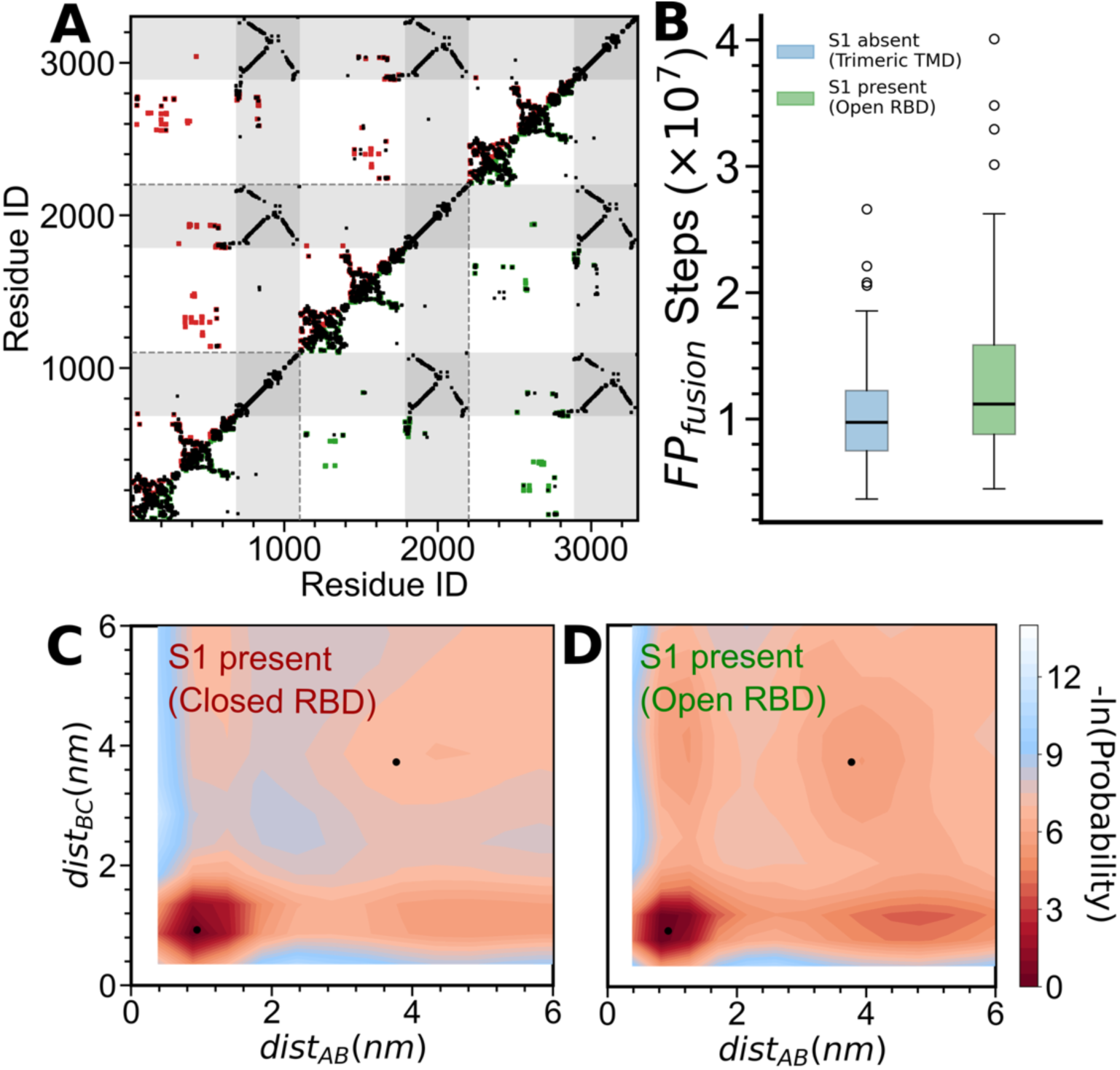
Coupling between the RBD opening and TMD dissociation. (A) The contact map used to build the S1-closed RBD (upper triangle) and S1-open RBD (lower triangle) models. A square shown at (i, j) indicates that a contact is present between residues i and j. Contacts common to both models (black) and closed-RBD (red) and open-RBD (green) model-specific contacts are shown. The threefold symmetry of the spike can be seen. The S2 subunits are marked by grey bands and are modeled using postfusion-state contacts, whereas the S1 subunit and inter-subunit contacts are taken from their respective prefusion structures. (B) Comparison of FP_fusion_, defined as the number of steps required for the average distance between the centers of mass of the three fusion peptides (d_FP_) to reach 1.27 nm shows that having the S1 present (open RBD; green) increases the time for the conformational transition as compared to when S1 is absent (blue). (C, D) Two-dimensional free-energy plots with dist_AB_ (distance between the centers of mass of TM helices A and B) and dist_BC_ as the two coordinates. Only those parts of the trajectories were analyzed where S1 remains associated with S2, i.e., Q_intersubunit_>0. (C) S1-closed RBD and (D) S1-open RBD models.

An initial set of 100 prefusion structures was sampled from simulations of a Cα-SBM built purely on the prefusion state (both S2 and S1) with all RBDs in the closed conformation. These were used to initiate 100 independent simulations of 20×10^7^ simulation steps with each of S1-closed RBD and S1-open RBD Cα-SBMs. The closed (**Figure S8A)** and open (**Figure S8B**) states of the RBD were first verified and were found to be consistent with the model used for the simulations. We next analyzed the fraction of intersubunit (S1-S2) contacts (Q_intersubunit_) over the simulation steps. For the S1-closed RBD Cα-SBM, Q_intersubunit_ indicates that 20-50% of contacts are retained throughout the simulation, except in five trajectories where the S1 subunit is shed after half the simulation time (10×10^7^ steps; **Figure S8C**). In contrast, for the open-RBD model, Q_intersubunit_ decreases to zero for all trajectories within the simulation time (**Figure S8D**). Thus, the opening of one RBD with the concomitant reduction of S1-S2 contacts results in a higher probability of S1 shedding.

The presence of the S1 subunit is also expected to delay the formation of the extended pre-hairpin intermediate of S2, in which the FPs are in their postfusion conformation, with the average of the distances between the centers-of-mass of the three FPs (d_FP_) being 1.27 nm. The time it takes for d_FP_ to converge to 1.27 nm, termed FP_fusion_, is on average greater for S1-open RBD Cα-SBM than it is in the trimeric TMD Cα-SBM (of S2) when S1 is absent (**Figure 4B**; see **Figure S9A** for data from other models). The additional steps required for S1 shedding when S1 is present increase FP_fusion_ and delay the overall S2 conformational conversion.

To investigate the effect of the RBD conformation further, we calculated the fraction of prefusion and postfusion contacts in the S2 subunits for individual simulation snapshots and plotted 2D free-energy surfaces for the S1-closed RBD and S1-open RBD models (**Figure S9B**). S2 rarely converts to the postfusion state in the S1-closed RBD Cα-SBM, whereas the S1-open RBD model undergoes a successful transition. Overall, these results indicate that closed RBDs inhibit S1 dissociation and spike transition, while opening a single RBD is sufficient to trigger these events.

Finally, we examine the effect of S1 presence and RBD conformation on the TMD population. The average interhelical distance in the TMD across simulations (**Figure S9C**) shows that the interhelical distance has large (unrestrained TMD) and small (trimeric TMD) fluctuations around the postfusion state when S1 is absent. In contrast, the average interhelical distance mostly stays below or at the postfusion state distance when S1 is present. However, the presence of S1 delays conversion to the post-fusion state and consequently to the post-fusion distance, with the S1-closed-RBD fluctuating up to the post-fusion state almost at the end of simulation time. To further quantify the differences in TMD dynamics for a given RBD conformation, we calculated the distances between the center-of-mass of helix A and helix B (dist_AB_), and helix B and helix C (dist_BC_) of the TMD from those sections of the trajectories when S1 has not yet been shed (Q_intersubunit_ > 0). The 2D free energy plot using dist_AB_ and dist_BC_ from S1-closed RBD trajectories (**Figure 4C**; compare with both **Figure 3D** and **3E**) shows that the TMD largely remains in the prefusion conformation, suggesting that S1 with closed RBDs increases the population of the TMD prefusion state and stabilizes it. Although the S1-open RBD also has a higher population of the prefusion TMD, the probability of sampling larger distances, specifically conformations with one helix dissociated from the prefusion TMD trimer, increases (**Figure 4D**). However, the presence of one open RBD eventually causes S1 to shed, and the TMD then transitions to the postfusion state (**Figure S9D**).

## DISCUSSION

### The role of viral fusion protein TMDs in membrane fusion

Although their size and membrane localization make it difficult to study the TMDs of viral and cellular proteins, it is known that they are important for membrane fusion (29). For instance, mutational studies of the vesicular stomatitis virus fusion protein TMD have shown that the substitution of glycine with alanine can abolish fusion (30). Similarly, replacing the influenza virus hemagglutinin TMD with a lipid anchor shows that the TMD amino acid sequence is required for stable fusion pore formation and subsequent fusion pore enlargement (31, 32). Recent simulations have highlighted the importance of intra-TMD and TMD-membrane interactions in membrane fusion (17, 33, 34). Here, we expand the possible role of the TMD in viral fusion by showing that it can modulate the timing of the conformational transition of the ectomembrane domains of viral fusion proteins.

### TMD dynamics can modulate the timing of the spike conformational transition

We first simulated the conformational conversion of the spike-S2 using a Cα-SBM, which encodes the postfusion structure of the SARS-CoV-2 spike-S2, and found that it was consistent with previous simulations (8, 11) (**Figure 2**). This postfusion structure had few intra-TMD contacts, which led to a dynamic TMD. We then performed the same simulations, but with a trimeric TMD stabilized by contacts derived from an NMR structure of the isolated spike-TMD. We find that constraining the TMD allows the fusion peptide (FP; **Figure 1**) to extend further and remain in the extended state for a longer duration, resulting in a slower conformational transition (**Figure 3**). We observe a correlation between the number of steps required for TMD opening and the steps required to complete the conformational transition (**Figure 3F**), supporting the notion that TMD dynamics could govern the kinetics of the conformational change.

If the TMD is overly dynamic, the S2 subunit may rapidly collapse into the postfusion conformation, potentially leading to postfusion structures that are nonproductive and do not bring about membrane fusion because the FPs fail to dock with the host membrane. Conversely, if the trimeric TMD is excessively constrained, spike-S2 may become kinetically trapped in late-fusion intermediate conformations, similar to those seen experimentally in the absence of S2’ cleavage (7). The timing of the S2 conformational transition is likely to be finely regulated and tuning the strength of intra-TMD interactions through sequence evolution may be one method to achieve this regulation.

Cholesterol is known to affect the free energy barriers associated with conformational changes in the membrane regions of pore-forming toxins by modulating their conformational flexibility (35, 36). Additionally, cholesterol treatment of both viral and host cell membranes reduces viral infectivity (37). A possible mechanism for this effect could be through cholesterol modulating TMD dynamics (17) and, in turn, affecting the fusion protein conformational transition.

### The presence of S1 stabilizes trimeric TMD

Simulations of the S2 conformational conversion with a trimeric TMD were performed in the presence of S1-closed RBD (S1 in the prefusion conformation and all RBDs in the closed conformation) and S1-open RBD (S1 in the prefusion conformation and one RBD in the open conformation), with the contacts between S1 and S2 being derived from the prefusion state. ACE2-bound RBD is expected to be in an open state (38), however, a dynamic RBD could also fluctuate between open and closed states (39, 40). We found that there was an 18% loss of S1-S2 contacts upon the opening of one RBD. This loss of contacts makes the S1-S2 interface less stable and amenable to breaking, which results in S1 shedding (**Figure S8**). In practice, even cleavage at the S1/S2 furin cleavage site has been shown to produce an inherently metastable state that is prone to S1 shedding even before ACE2 engagement (1, 26).

Overall, the population of trimeric TMD states is increased in the presence of S1 (both open RBD and closed RBD) (**Figure 4C** and **4D**) as compared with its absence (**Figure S9D**). Additionally, the S1-open RBD model has a higher population of non-trimeric TMD states than the S1-closed RBD model (**Figure 4D**). This higher dissociated TMD population could amplify the effect of S1-S2 contact loss and drive a faster conformational conversion of S2 to its postfusion state.

### Modelling assumptions and limitations

The present Cα-SBM makes two main assumptions: First, the prefusion structure of S2 is unstable and locked into place via the S1 cap. Frustration calculations (41) show that the postfusion state of S2 is less frustrated than its prefusion state (**Figure S2B**), providing support for the validity of this assumption. Additionally, it is known that some percentage of spike proteins shed S1 and are present in the postfusion conformation in the viral membrane (26), further supporting this assumption. In practice, at least some intra-S2 prefusion structure interactions are likely to be stable, and these will modulate the population of the various intermediates and the timing of the conformational conversion. However, the Cα-SBM is a physically reasonable and computationally efficient starting point for understanding the effect of TMD dynamics upon the S2 conformational transition, which is the aim of the present study.

Secondly, an implicit representation of the membrane is used, which keeps the TMD anchored but does not modulate its dynamics through specific membrane-protein interactions. SBM simulations investigate how broad structural features of the protein contribute to conformational transition mechanisms. By introducing two distinct TMDs, one of which is unrestrained, while the other has a stable trimeric structural ensemble, we investigate whether TMD dynamics can influence the mechanism of conformational conversion. The two distinct types of TMD can arise in practice due to differences in the length and nature of the TMD sequences (42), the types of lipids present in the membrane, etc.

We have also introduced the two different conformations of S1 (all closed RBD and one open RBD) independently into the simulations, rather than developing a dual-SBM for S1 (43). In the present simulations, we find that the presence of S1 increases the population of the trimeric TMD. However, it is possible that the TMD conformation itself, specifically the non-trimeric dissociated TMD, could promote RBD opening for easy access to ACE2. Whether this is a possibility can be explored through the development of a dual-SBM for S1. It should also be noted that because of the unstable prefusion S2 conformation, S2 can start converting to the postfusion state while S1 remains attached (or has not shed), especially in the S1-open RBD model. Whether this occurs in experiments may depend on the details of the energetics of various terms, which the present coarse-grained model does not include.

Finally, in the present model, S1 (both closed and open RBD models) sheds as a trimer, and the FPs zip up into a fusion head cooperatively. Although these processes are diverse, we assess them together because they assume structure-derived strong inter-protomer interactions within the spike. There is some evidence in the SARS-CoV-2 spike (12, 44) as well as in HIV (45) that the inter-protomer interfaces breathe, which may lead to a reduced cooperativity in dynamics not captured in the present Cα-SBMs. However, a recently resolved structure (12) indicates that the fusion head may form cooperatively.

### The TMD as a potential target for fusion inhibitors

Spike S2-TMD dynamics affect the residence time of the I1 intermediate (**Figure 3C**; akin to the extended prehairpin intermediate seen experimentally (7–10)) and thus, the amount of time the FPs have to find and dock with the host membrane (**Figure 1**) to drive viral fusion. The previous all atom SBM simulations (11) showed that glycans decorating S2 could increase the population of I1-like intermediates and help the fusion head dock with the host membrane. Here, we show that modulating TMD dynamics can also alter the population of I1 and could, in turn, be another method for therapeutic intervention. One way to modulate TMD dynamics could be to use small lipid-like molecules that bind the TMD but also change local membrane fluidity. In fact, host-derived antiviral peptides such as IFITMs modulate local rigidity through lipid sorting and can block enveloped virus fusion, highlighting the feasibility of this strategy (46, 47). Recent bioinformatic analyses show that, on average, class I viral fusion proteins possess longer TMD helices than human fusion proteins, a feature that may facilitate TMD dynamics (42). Thus, non-specific lipid-like modulators of TMD dynamics could function as broad-spectrum inhibitors of class I viral fusion proteins.

## CONCLUSIONS

The SARS-CoV-2 spike protein is composed of two parts: S1, the host recognition subunit, which caps S2, the fusion subunit. S2 undergoes a large conformational transition, from a prefusion to a postfusion structure, which facilitates membrane fusion. In its prefusion state, S2 is anchored to the viral membrane through three single-pass transmembrane helices, together known as the transmembrane domain (TMD), and often regarded as a passive trimer. However, mutational studies indicate that the TMD may contribute to SARS-CoV-2 function and membrane fusion. Here, we use molecular dynamics simulations of a coarse-grained structure-based model of S2 to study its conformational conversion from the prefusion to the postfusion state. We find that two intermediates are populated during this conformational transition, whose structures are similar to recently experimentally resolved conformations of the spike protein. Increased TMD dynamics and TMD dissociation reduce the population of these intermediates and increase the rate of the S2 conformational transition. Despite not being directly in contact with the TMD, the presence of the S1 cap arrests S2 in its prefusion conformation, reduces TMD dynamics, and delays the S2 conformational transition. However, a receptor-bound conformation of the S1 cap can ease the constraints on the TMD and allow it to be more dynamic. Since local membrane fluidity is also likely to affect TMD dynamics, the TMD may be able to integrate signals from both its membrane environment and the conformational state of the spike-S1 and tune the timing of the S2 conformational transition. Conversely, molecules that alter TMD dynamics may modulate the S2 conformational conversion and offer new avenues for antiviral strategies. Overall, despite being a small and understudied part of the spike, the TMD may contribute substantially to its function.

## METHODS

### Model rationalization and methodology

The spike-S2 conformational transition is triggered upon S2’ cleavage and S1 subunit dissociation (1). Therefore, the S2 subunit can be treated as a loaded spring in the prefusion state, which is released upon S1 dissociation. The frustratometer (41), which quantifies energetic frustration (and in turn destabilization) in protein structures, also shows that the postfusion structure is less frustrated and therefore more stable than the prefusion state (also see **Figure S2B**). Based on these observations, the previous all-atom SBM simulations of the spike-S2 (11) were initiated from the pre-fusion structure of spike-S2 while using an SBM based purely on the post-fusion structure of spike-S2. Consequently, a conversion from the prefusion state to the postfusion state was seen, driven by the potential energy function. Here, we follow the same reasoning and methodology, but using a recently resolved post-fusion structure (16).

### Protein structures

Most of the postfusion structure (PDB ID: 8FDW; missing residues: parts of the S2-S2’ linker domain, residues 686-694 and 785-831, and the CT domain, residues 1250-1273) (16) has been resolved. The present Cα-SBM of S2 used the residues 834 to 1249 of the postfusion structure, which starts at the S2’ cleavage site and contains all CT domain residues which have been resolved and contains: the fusion peptide (FP), heptad repeats (HR1 and HR2), the transmembrane domain (TMD), and a small part of the cytoplasmic tail (see **Figure 1** and **Figure 2A**). The prefusion structure with all three RBDs in the closed state was taken from PDB ID: 6VXX (48), and the structure with a single RBD in the open state was taken from PDB ID: 6VSB (38). These structures were not fully resolved. However, complete modeled structures are available from the CHARMM-GUI COVID-19 Archive (49, 50). The structures used in the simulations have been deposited as supplementary data.

The prefusion TMD trimer structure was obtained from the NMR-resolved structure (PDB ID 7LC8; residues 1217-1237), which showed that the isolated TMD exists as a trimer in bicelles (28). Note that the structure contains 5 mutations (M1229L, M1233L, C1235S, C1236S, and M1237T), with the sequence WLGFIAGLIAIVLVTILLSST.

### Cα-Structure-Based Model

We used a common form of the Cα-SBM (51) to build the potential energy function (E):

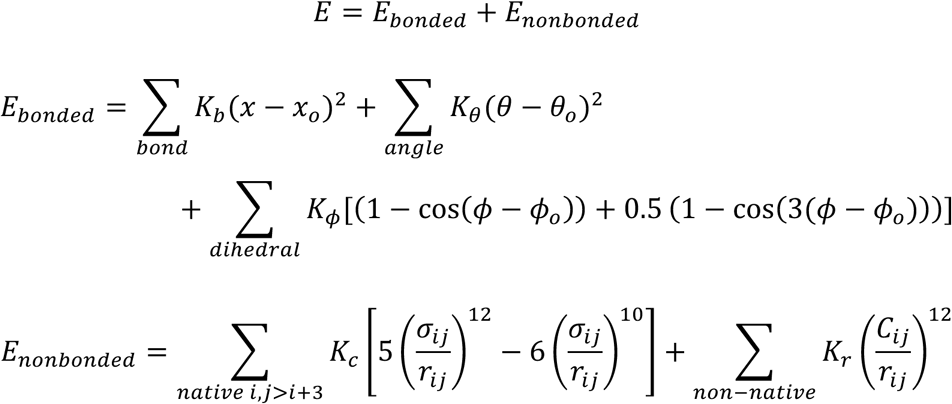

where x_o_, θ_o_, and ϕ_o_ are the native values (calculated from the structure on which the Cα-SBM is built) of bond lengths, angles, and dihedrals, respectively. Bond and angles are modeled using a harmonic potential, whereas the dihedrals are modeled using a combination of cosine functions. The strength of individual interaction potentials is set to K_b_ = 100ε, K_θ_ = 20ε, and K_ϕ_ = 1ε, where ε is the reference energy scale, whose value here is set to 1 kJ/mol. Non-bonded interactions are defined between pairs of residues (i, j) separated by more than 3 residues in sequence. Attractive 10-12 Lennard-Jones (LJ)-like interactions are defined between those residue pairs that are in contact in the native structure. The minima (σ_ij_) are located at the native distance between the two Cα atoms defined to be in contact. The interaction strength of this LJ-like potential is set to K_c_=1ε for all the contact pairs. A contact is present between residues i and j if at least one heavy atom of i is within 0.45 nm of at least one heavy atom of j. A repulsive potential is present between pairs of residues not in contact, with the strength K_r_ set to 1ε and C_ij_=0.4 nm.

### Implicit membrane potential

A “reverse flat-bottom potential” was used to keep the membrane-flanking residues (HG and HR2 on one side and CT on the other, **Figure 1**) from entering the implicit membrane region, which was present along the Z-coordinate plane. The potential is defined as:

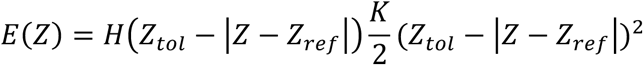

where E(Z) is the potential energy as a function of the Z-coordinate of the atoms, Z_ref_ is the center of the implicit membrane (defined as Z-coordinate of the center-of-mass of the TMD: residues 1211–1234), Z_tol_ defines the implicit membrane region around the membrane center (2.5 nm on each side), and K is the harmonic potential strength set to 10ε/nm^2^. H is the rectangular window function implemented using the Heaviside step function. E(Z) with Z_ref_=0 is plotted in **Figure S1**. The potential was applied to the head region HG (residues 1068-1128), HR2 region (residues 1162-1208), and CT region (residues 1247-1249). Since the transition begins with the head moving towards the membrane (11), the HG region was selected to prevent a head-membrane interaction. The HR2 and CT regions were selected to maintain the TMD within the implicit membrane region and to prevent HR2 and CT from inserting into the membrane. Selecting the membrane-flanking regions HR2 and CT, automatically keeps the TMD in the membrane region without the need for an additional potential.

### Simulation Models

The Unrestrained TMD model was a Cα-SBM built solely on the postfusion structure of S2 (FP to CT: residues 834-1249). For the Trimeric TMD model, intra-TMD contacts are first calculated from the trimeric TMD structure (PDB ID 7LC8), and these 111 contacts (presumed to be present in the prefusion state of S2) are added to the Unrestrained TMD model. These contacts had a K_c_=0.8ε to avoid perturbing the final postfusion state (see Results and **Figure S4**).

For the S1/S2 models, S1 (residues 1-685) was modeled, and the S1-S2 contacts were calculated using prefusion structures with either all RBDs closed (S1-closed RBD model) or with one RBD open (S1-open RBD model). For both models, S2 (834–1249) was modeled using the trimeric TMD model. All relevant structures and contact maps used in the study are deposited as supplementary data.

### Simulation protocols

All input files for the Cα-SBMs were generated using the SMOG 2 software package (52) with additional in-house scripts used to modify the force field for different model variants. All simulations were performed using the OpenSMOG software package (v1.2) (53) with additional code modifications to implement the implicit membrane potential.

The temperature scale in SBMs is measured in reduced units of k_B_T, with k_B_ = 0.00831451. To identify the optimal temperature for simulating the transition within a feasible simulation time, we first determined the highest reduced temperature at which the protein can transition to a stable postfusion state. We performed 10 independent transition simulations of 2× 10^8^ steps starting from the S2 prefusion state using the Cα-SBM of the S2 Unrestrained TMD model and quantified the fraction of formed native-contacts (Q) for both the entire protein (2410 contacts) and the TMD (99 contacts) in the postfusion ensemble. As shown in **Figure S3A**, increasing the reduced temperature from 0.67 to 0.75 leads to a slight decrease in protein contacts from Q ≍ 0.94 to Q ≍ 0.90, while TMD contacts decrease from Q ≍ 0.75 to Q ≍ 0.65. At 0.79, the protein contacts drop further to Q ≍ 0.88 with minimal change in the TMD contacts. At 0.83, both Q values continue to decrease and exhibit substantial variation in the mean estimates, indicating partial destabilization. At 0.87, the protein and TMD undergo substantial unfolding and loss of native structure. Based on these results, a reduced temperature of 0.75 maintains both the protein and the TMD in a stable state after fusion and allows the transition to occur within a feasible simulation time. We performed all subsequent transition simulations at this reduced temperature.

To generate initial prefusion conformations, we built two Cα-SBMs based on (i) the prefusion structure of S2 with simulation initiated from the same structure and (ii) the prefusion structure of the S1/S2 complex with S1 having all RBDs closed with simulation initiated from the same structure. One simulation of both these Cα-SBMs was performed at 0.41 reduced temperature, and 100 conformations were randomly sampled from each simulation to generate initial prefusion structures. These structures were subsequently used to initiate conformational conversion simulations.

A total of four sets of simulations were performed, all initiated from prefusion structures. Each simulation was first energy-minimized using the L-BFGS minimizer with a maximum force tolerance of 1 kJ/mol/nm. The transition simulation runs were performed using the Langevin dynamics integrator with a time step of 0.5 ps, collision rate of 1.0 ps^-1^, non-bonded interaction cutoff of 3.0 nm, and a reduced temperature of 0.75. All simulations were run for a total of 2 × 10^8^ steps, and trajectories were saved every 2000 steps. A complete list of the performed simulations is given in **Table S1**. The conformational conversion simulations were performed with 10 non-interacting replicas placed in a box to generate 10 replicates on a single GPU. The input files for the non-interacting replicas were generated using SuBMIT (Structure Based Model(s) Input Toolkit; https://github.com/sglabncbs/submit). This setup allows us to use the available GPUs efficiently. A comprehensive description of this method is currently in preparation.

### Simulation analyses

The following structural parameters were used to characterize the conformational changes in the SARS-CoV-2 spike protein:

1. Center of Mass (com) of a specific region (e.g. FP), is calculated by using all residues in that region (residues 834–910 for the FP) from all three chains A, B, and C of the spike homotrimer. When com is only for one chain, this is explicitly stated.
2. Fraction of formed native contacts of a given type, X (Q_X_): A contact is said to be formed if the distance between the pair of residues that form it is less than 1.2 times their native distance. The number of such formed contacts (of type X; can be all) in a simulation snapshot is counted and divided by the total number of native contacts (of type X) to get the Q_X_ for that snapshot.
3. Root Mean Square Deviation (RMSD): RMSD of a simulation snapshot was computed relative to the postfusion state to track the conformational change. RMSD values near zero indicate a close to complete conversion to the postfusion structure.
4. FP_Z_ is defined as

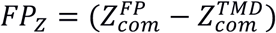

where 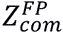 is the Z coordinate of the FP (residues 834–910) center of mass, and 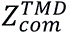 is the Z-coordinate of the TMD (residues 1211–1234) center of mass. The FP_Z_ values for the prefusion and postfusion states are 17.5 and 0.67 nm, respectively. Since the TMD helices are constrained within the implicit membrane region, FP_Z_ is a readout of the FP dynamics in the direction perpendicular to the membrane.
5. FP_completion_ is the number of simulation steps it takes for the FP to convert into a postfusion like structure, defined a FP_Z_ < 0.67 nm.
6. Z_head_ and r_head_: These parameters quantify the global movement of the head (HG; residues 1068-1128) relative to the TMD (residues 1211-1234, chains), and are defined as

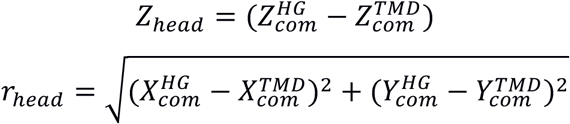

where (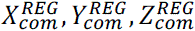) is the position of the center of mass (com) of REG (HG or TMD). *Z*_*head*_, the difference between the Z-coordinates, represents the vertical separation between the TMD and HG. *r*_*head*_ represents the lateral (radial) distance in the XY-plane between the TMD and the HG. The (Z_head_, r_head_) values are (15.3 nm, 0.025 nm) for the prefusion and (21.8 nm, 0.015 nm) for the postfusion native structures.
7. dist_AB_, dist_BC_, and dist_CA_: These are the distances between the centers of mass (COM) of the transmembrane domain (residues 1211–1234) of chains A and B (dist_AB_), B and C (dist_BC_), and C and A (dist_CA_) (see **Figure S7A**). These distances are used to quantify internal structural reorganization within the TMD during the transition from the prefusion to postfusion states.
8. 2D free energy surfaces (2DFES) or plots (2DFEPs) were computed by binning the number of simulation snapshots using two structural coordinates (such as dist_AB_ and dist_BC_) or calculating the 2D joint probabilities across all replicates, followed by computing the -ln(probability). Since we perform transition simulations, this is not a true “free energy” and is only analogous to that in equilibrium simulations. For the (Z_head_, r_head_) 2DFES calculation, the trajectories were truncated close to the point at which they converged to the postfusion state to avoid oversampling the final postfusion state. Trajectories were truncated when the values of (Z_head_, r_head_) matched the windows (21.5 nm<Z_head_<22.5 nm, 2 nm<r_head_<10 nm). Also, see **Figure S6A**.
9. Average interhelical distance is defined as the average of dist_AB_, dist_BC_, and dist_CA,_ and its value is 0.93 nm in the prefusion state and 3.5 nm in the postfusion state. The interhelical distance was also averaged over the 100 replicates to understand the nature of TMD dynamics across the different models.
10. TM_opening_ is defined as the number of steps it takes for the TM domain to dissociate or open. The TM domain is considered to be open when the interhelical distance exceeds that in the prefusion state (0.93 nm) and remains above it.
11. RBD angles: Residues 445 from the RBDs of chains A, B, and C form a triangle (**Figure S8A** inset). The angles at each triangle vertex were calculated from all simulation snapshots to track RBD opening (see **Figure S8A** and **S8B**).
12. FP_fusion_ is defined as the number of steps required for the average distance between pairs of centers of mass of the FPs of chains A, B, C (d_FP_) to reach 1.27 nm. This provides a measure of the steps required for FP association.

## Author contributions

AK, AB, SL, and SG designed research; AB and SL performed preliminary simulations; AK tuned the model, performed production simulations and analysis; AK wrote the first draft; AK, SL, AB, and SG updated the draft.

## Acknowledgments

This work was funded by the Department of Atomic Energy, Government of India through the Tata Institute of Fundamental Research (Project Identification No. RTI 4006). SG was supported by a grant from the National Supercomputing Mission (NSM) through the grant MeitY/R\&D/HPC/2(1)/2014 and SERB Grant: CRG/2021/004754. AK would like to thank the ANRF for providing the National Postdoctoral Fellowship (PDF/2025/002723) to pursue this research project. We thank Dr. Digvijay Lalwani Prakash, National Centre for Biological Sciences, Bangalore, for help in setting up the non-interacting replicas.

## Data Availability

Information required to reproduce the simulation data using the open-source OpenSMOG program is provided in the manuscript or included in the Supplementary Information. The full codes and input files will be made available upon publication of the manuscript.

## References

1. Jackson, C.B., M. Farzan, B. Chen, and H. Choe. 2022. Mechanisms of SARS-CoV-2 entry into cells. Nat. Rev. Mol. Cell Biol. 23:3–20.

2. Rossi, G.P., P. Anand, A. Puranik, M. Aravamudan, A.J. Venkatakrishnan, and V. Soundararajan. 2020. SARS-CoV-2 strategically mimics proteolytic activation of human ENaC. Elife 9:e58603.

3. Andersen, K.G., A. Rambaut, W.I. Lipkin, E.C. Holmes, and R.F. Garry. 2020. The proximal origin of SARS-CoV-2. Nat. Med. 26:450–452.

4. Shang, J., Y. Wan, C. Luo, G. Ye, Q. Geng, A. Auerbach, and F. Li. 2020. Cell entry mechanisms of SARS-CoV-2. Proc. Natl. Acad. Sci. U. S. A. 117:11727–11734.

5. Rencilin, C.F., A. Chatterjee, M.Y. Ansari, S. Deshpande, S. Mukherjee, R. Singh, S.B. Jayatheertha, P.M. Reddy, N. Hingankar, R. Varadarajan, J. Bhattacharya, and S. Dutta. 2025. Cryo-EM reveals conformational variability in the SARS-CoV-2 spike protein RBD induced by two broadly neutralizing monoclonal antibodies. RSC Adv. 15:14385–14399.

6. Saha, P., I. Fernandez, F. Sumbul, C. Valotteau, D. Kostrz, A. Meola, E. Baquero, A. Sharma, J.R. Portman, F. Stransky, T. Boudier, P. Guardado-Calvo, C. Gosse, T. Strick, F.A. Rey, and F. Rico. 2025. Modulation of SARS-CoV-2 spike binding to ACE2 through conformational selection. Nat. Nanotechnol. 20:926–934.

7. Akıl, C., J. Xu, J. Shen, and P. Zhang. 2025. Unveiling the structural spectrum of SARS-CoV-2 fusion by in situ cryo-ET. Nat. Commun. 16:1–12.

8. Grunst, M.W., Z. Qin, E. Dodero-Rojas, S. Ding, J. Prévost, Y. Chen, Y. Hu, M. Pazgier, S. Wu, X. Xie, A. Finzi, J.N. Onuchic, P.C. Whitford, W. Mothes, and W. Li. 2024. Structure and inhibition of SARS-CoV-2 spike refolding in membranes. Science 385:757–765.

9. Su, R., J. Zeng, T.C. Marcink, M. Porotto, A. Moscona, and B. O’Shaughnessy. 2023. Host Cell Membrane Capture by the SARS-CoV-2 Spike Protein Fusion Intermediate. ACS Cent. Sci. 9:1213–1228.

10. Marcink, T.C., T. Kicmal, E. Armbruster, Z. Zhang, G. Zipursky, K.L. Golub, M. Idris, J. Khao, J. Drew-Bear, G. McGill, T. Gallagher, M. Porotto, A. Des Georges, and A. Moscona. 2022. Intermediates in SARS-CoV-2 spike– mediated cell entry. Sci. Adv. 8:eabo3153.

11. Dodero-Rojas, E., J.N. Onuchic, and P.C. Whitford. 2021. Sterically confined rearrangements of SARS-CoV-2 Spike protein control cell invasion. Elife 10:e70362.

12. Xing, L., Z. Liu, X. Wang, Q. Liu, W. Xu, Q. Mao, X. Zhang, A. Hao, S. Xia, Z. Liu, L. Sun, G. Zhang, Q. Wang, Z. Chen, S. Jiang, L. Sun, and L. Lu. 2025. Early fusion intermediate of ACE2-using coronavirus spike acting as an antiviral target. Cell 188:1297–1314.e24.

13. Lu, Y., T.L. Neo, D.X. Liu, and J.P. Tam. 2008. Importance of SARS-CoV spike protein Trp-rich region in viral infectivity. Biochem. Biophys. Res. Commun. 371:356–360.

14. Corver, J., R. Broer, P. Van Kasteren, and W. Spaan. 2009. Mutagenesis of the transmembrane domain of the SARS coronavirus spike glycoprotein: Refinement of the requirements for SARS coronavirus cell entry. Virol. J. 6:230.

15. Ortiz-Mateu, J., D. Belda, A.I. Avilés-Alía, J. Alonso-Romero, M.J. García-Murria, I. Mingarro, R. Geller, and L. Martinez-Gil. 2025. The sequence and structural integrity of the SARS-CoV-2 Spike protein transmembrane domain is crucial for viral entry. *Commun*. Biol. 8:1579.

16. Shi, W., Y. Cai, H. Zhu, H. Peng, J. Voyer, S. Rits-Volloch, H. Cao, M.L. Mayer, K. Song, C. Xu, J. Lu, J. Zhang, and B. Chen. 2023. Cryo-EM structure of SARS-CoV-2 postfusion spike in membrane. Nature 619:403–409.

17. Lall, S., P. Balaram, M.K. Mathew, and S. Gosavi. 2025. Sequence of the SARS-CoV-2 Spike Transmembrane Domain Encodes Conformational Dynamics. J. Phys. Chem. B 129:194–209.

18. Bryngelson, J.D., J.N. Onuchic, N.D. Socci, and P.G. Wolynes. 1995. Funnels, pathways, and the energy landscape of protein folding: a synthesis. Proteins Struct. Funct. Bioinforma. 21:167–195.

19. Chavez, L.L., J.N. Onuchic, and C. Clementi. 2004. Quantifying the roughness on the free energy landscape: entropic bottlenecks and protein folding rates. J. Am. Chem. Soc. 126:8426–8432.

20. Whitford, P.C., J.K. Noel, S. Gosavi, A. Schug, K.Y. Sanbonmatsu, and J.N. Onuchic. 2009. An all-atom structure-based potential for proteins: bridging minimal models with all-atom empirical forcefields. Proteins Struct. Funct. Bioinforma. 75:430–441.

21. Kulshrestha, A., T.M. Phan, A. Rizuan, P. Mohanty, and J. Mittal. 2025. Multiscale Simulations Elucidate the Mechanism of Polyglutamine Aggregation and the Role of Flanking Domains in Fibril Polymorphism. J. Phys. Chem. B 129:11205–11219.

22. Giri Rao, V.V.H., R. Desikan, K.G. Ayappa, and S. Gosavi. 2016. Capturing the membrane-triggered conformational transition of an α-helical pore-forming toxin. J. Phys. Chem. B 120:12064–12078.

23. Kulshrestha, A., S.N. Punnathanam, and K.G. Ayappa. 2022. Finite temperature string method with umbrella sampling using path collective variables: application to secondary structure change in a protein. Soft Matter 7593–7603.

24. Shi, W., G.M. Jonaid, M.G. Kibria, J. Allen, H. Peng, S. Rits-Volloch, H. Zhu, S. Wang, R.M. Walsh, J. Lu, and B. Chen. 2025. Effect of the S2’ site cleavage on SARS-CoV-2 spike. Nat. Commun. 16:11675.

25. Benton, D.J., A.G. Wrobel, P. Xu, C. Roustan, S.R. Martin, P.B. Rosenthal, J.J. Skehel, and S.J. Gamblin. 2020. Receptor binding and priming of the spike protein of SARS-CoV-2 for membrane fusion. Nature 588:327– 330.

26. Ke, Z., J. Oton, K. Qu, M. Cortese, V. Zila, L. McKeane, T. Nakane, J. Zivanov, C.J. Neufeldt, B. Cerikan, J.M. Lu, J. Peukes, X. Xiong, H.G. Kräusslich, S.H.W. Scheres, R. Bartenschlager, and J.A.G. Briggs. 2020. Structures and distributions of SARS-CoV-2 spike proteins on intact virions. Nature 588:498–502.

27. Eddy, N.R., and J.N. Onuchic. 2018. Rotation-Activated and Cooperative Zipping Characterize Class I Viral Fusion Protein Dynamics. Biophys. J. 114:1878–1888.

28. Fu, Q., and J.J. Chou. 2021. A Trimeric Hydrophobic Zipper Mediates the Intramembrane Assembly of SARS-CoV-2 Spike. J. Am. Chem. Soc. 143:8543–8546.

29. Langosch, D., M. Hofmann, and C. Ungermann. 2007. The role of transmembrane domains in membrane fusion. Cell. Mol. Life Sci. 64:850–864.

30. Cleverley, D.Z., and J. Lenard. 1998. The transmembrane domain in viral fusion: Essential role for a conserved glycine residue in vesicular stomatitis virus G protein. Proc. Natl. Acad. Sci. U. S. A. 95:3425–3430.

31. Markosyan, R.M., F.S. Cohen, and G.B. Melikyan. 2000. The lipid-anchored ectodomain of influenza virus hemagglutinin (GPI-HA) is capable of inducing nonenlarging fusion pores. Mol. Biol. Cell 11:1143–1152.

32. Kemble, G.W., T. Danieli, and J.M. White. 1994. Lipid-anchored influenza hemagglutinin promotes hemifusion, not complete fusion. Cell 76:383–391.

33. van Tilburg, M., P.A.J. Hilbers, and A.J. Markvoort. 2023. On the role of membrane embedding, protein rigidity and transmembrane length in lipid membrane fusion. Soft Matter 19:1791–1802.

34. Scherer, K.C., C.S. Poojari, and J.S. Hub. 2026. Transmembrane domains of fusion proteins promote stalk formation by inducing membrane disorder. Biophys. J. 125:1276–1285.

35. Kulshrestha, A., S.N. Punnathanam, R. Roy, and K.G. Ayappa. 2023. Cholesterol catalyzes unfolding in membrane-inserted motifs of the pore forming protein cytolysin A. Biophys. J. 122:4068–4081.

36. Kulshrestha, A., S. Maurya, T. Gupta, R. Roy, S.N. Punnathanam, and K.G. Ayappa. 2023. Conformational Flexibility Is a Key Determinant for the Lytic Activity of the Pore-Forming Protein, Cytolysin A. J. Phys. Chem. B 127:69–84.

37. Sanders, D.W., C.C. Jumper, P.J. Ackerman, D. Bracha, A. Donlic, H. Kim, D. Kenney, I. Castello-Serrano, S. Suzuki, T. Tamura, A.H. Tavares, M. Saeed, A.S. Holehouse, A. Ploss, I. Levental, F. Douam, R.F. Padera, B.D. Levy, and C.P. Brangwynne. 2021. SARS-CoV-2 requires cholesterol for viral entry and pathological syncytia formation. Elife 10:e65962.

38. Wrapp, D., N. Wang, K.S. Corbett, J.A. Goldsmith, C.-L. Hsieh, O. Abiona, B.S. Graham, and J.S. McLellan. 2020. Cryo-EM structure of the 2019-nCoV spike in the prefusion conformation. Science (80*-.).* 367:1260– 1263.

39. Gur, M., E. Taka, S.Z. Yilmaz, C. Kilinc, U. Aktas, and M. Golcuk. 2020. Conformational transition of SARS-CoV-2 spike glycoprotein between its closed and open states. J. Chem. Phys. 153.

40. Berger, I., and C. Schaffitzel. 2020. The SARS-CoV-2 spike protein: balancing stability and infectivity. Cell Res. 30:1059–1060.

41. Parra, R.G., N.P. Schafer, L.G. Radusky, M.Y. Tsai, A.B. Guzovsky, P.G. Wolynes, and D.U. Ferreiro. 2016. Protein Frustratometer 2: a tool to localize energetic frustration in protein molecules, now with electrostatics. Nucleic Acids Res. 44:W356–W360.

42. Lall, S., P. Balaram, M.K. Mathew, and S. Gosavi. 2025. The longer transmembrane helices of class I viral fusion proteins may facilitate viral fusion. bioRxiv 2025.09.25.678357.

43. Whitford, P.C., O. Miyashita, Y. Levy, and J.N. Onuchic. 2007. Conformational Transitions of Adenylate Kinase: Switching by Cracking. J. Mol. Biol. 366:1661–1671.

44. Costello, S.M., S.R. Shoemaker, H.T. Hobbs, A.W. Nguyen, C.L. Hsieh, J.A. Maynard, J.S. McLellan, J.E. Pak, and S. Marqusee. 2022. The SARS-CoV-2 spike reversibly samples an open-trimer conformation exposing novel epitopes. Nat. Struct. Mol. Biol. 29:229–238.

45. Li, W., Z. Qin, E. Nand, M.W. Grunst, J.R. Grover, J.W. Bess, J.D. Lifson, M.B. Zwick, H.D. Tagare, P.D. Uchil, and W. Mothes. 2023. HIV-1 Env trimers asymmetrically engage CD4 receptors in membranes. Nature 623:1026–1033.

46. Klein, S., G. Golani, F. Lolicato, C. Lahr, D. Beyer, A. Herrmann, M. Wachsmuth-Melm, N. Reddmann, R. Brecht, M. Hosseinzadeh, A. Kolovou, J. Makroczyova, S. Peterl, M. Schorb, Y. Schwab, B. Brügger, W. Nickel, U.S. Schwarz, and P. Chlanda. 2023. IFITM3 blocks influenza virus entry by sorting lipids and stabilizing hemifusion. Cell Host Microbe 31:616–633.e20.

47. Shi, G., A.D. Kenney, E. Kudryashova, A. Zani, L. Zhang, K.K. Lai, L. Hall-Stoodley, R.T. Robinson, D.S. Kudryashov, A.A. Compton, and J.S. Yount. 2020. Opposing activities of IFITM proteins in SARS-CoV-2 infection. EMBO J. 2020 *403* 40:EMBJ2020106501.

48. Walls, A.C., Y.J. Park, M.A. Tortorici, A. Wall, A.T. McGuire, and D. Veesler. 2020. Structure, Function, and Antigenicity of the SARS-CoV-2 Spike Glycoprotein. Cell 181:281–292.e6.

49. Woo, H., S.J. Park, Y.K. Choi, T. Park, M. Tanveer, Y. Cao, N.R. Kern, J. Lee, M.S. Yeom, T.I. Croll, C. Seok, and W. Im. 2020. Developing a fully glycosylated full-length SARS-COV-2 spike protein model in a viral membrane. J. Phys. Chem. B 124:7128–7137.

50. Choi, Y.K., Y. Cao, M. Frank, H. Woo, S.J. Park, M.S. Yeom, T.I. Croll, C. Seok, and W. Im. 2021. Structure, Dynamics, Receptor Binding, and Antibody Binding of the Fully Glycosylated Full-Length SARS-CoV-2 Spike Protein in a Viral Membrane. J. Chem. Theory Comput. 17:2479–2487.

51. Clementi, C., H. Nymeyer, and J.N. Onuchic. 2000. Topological and energetic factors: what determines the structural details of the transition state ensemble and “en-route” intermediates for protein folding? an investigation for small globular proteins. J. Mol. Biol. 298:937–953.

52. Noel, J.K., M. Levi, M. Raghunathan, H. Lammert, R.L. Hayes, J.N. Onuchic, and P.C. Whitford. 2016. SMOG 2: a versatile software package for generating structure-based models. PLOS Comput. Biol. 12:e1004794.

53. de Oliveira, A.B., V.G. Contessoto, A. Hassan, S. Byju, A. Wang, Y. Wang, E. Dodero-Rojas, U. Mohanty, J.K. Noel, J.N. Onuchic, and P.C. Whitford. 2022. SMOG 2 and OpenSMOG: Extending the limits of structure-based models. Protein Sci. 31:158–172.

