## Supplementary Information for "The transmembrane domain regulates the kinetics of the SARS-CoV-2 spike conformational transition"

**for**

**Table S1: List of simulations.**

| | Initial state | SBM potential from | Number of steps | Replicates | Simulation |
|---|---|---|---|---|---|
| 1. | S2-prefusion | S2-prefusion | $2 \times 10^8$ | 1 | Equilibrium |
| 2. | S2-prefusion | Unrestrained TMD: S2-postfusion | $2 \times 10^8$ | 100 | Transition |
| 3. | S2-prefusion | Trimeric TMD: S2-postfusion+NMR derived contacts for the TMD | $2 \times 10^8$ | 100 | Transition |
| 4. | S1/S2 complex with closed RBD | S1-prefusion-closed RBD<br><br>S2-prefusion | $2 \times 10^8$ | 1 | Equilibrium |
| 5. | S1/S2 complex with closed RBD | S1-prefusion-closed RBD<br><br>S2-postfusion+NMR derived contacts for the TMD | $2 \times 10^8$ | 100 | Transition |
| 6. | S1/S2 complex with closed RBD | S1-prefusion-open RBD<br><br>S2-postfusion+NMR derived contacts for the TMD | $2 \times 10^8$ | 100 | Transition |

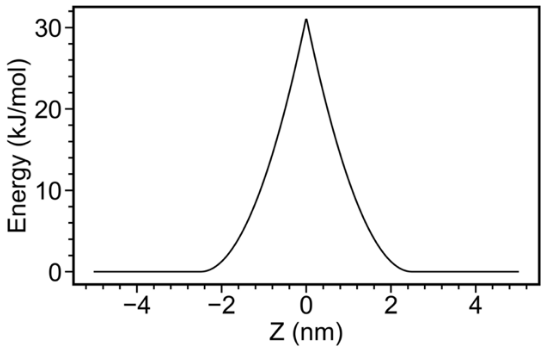


**Figure S1. Implicit membrane potential.** The TMD helices are confined to an implicit membrane region by applying a potential to regions flanking the TMD (HG and HR2 on one side and CT on the other). The functional form of this "reverse flat-bottom potential" is given in the Methods Section. Here, the membrane center is at $Z_{ref}=0$, and there is a tolerance region, $Z_{tol}=2.5$ nm, on both sides of the membrane center, resulting in an effective membrane thickness of 5 nm. A spring constant of 10 kJ/mol/nm$^2$ is used to enforce the restraint.

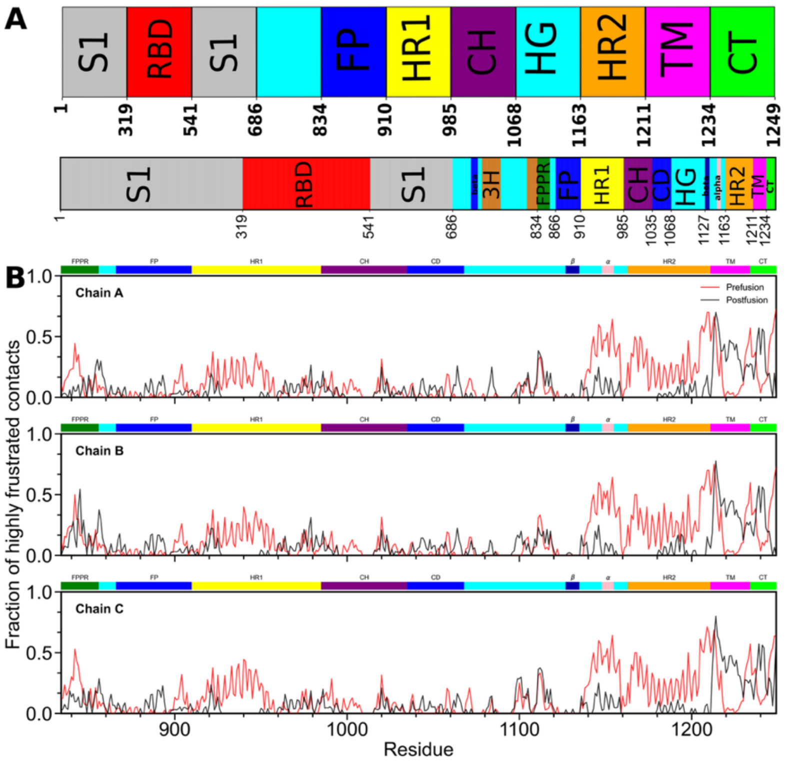


**Figure S2: Domain definitions of S and frustration in the S2 subunit.** (A) The coarse-grained domain definitions used here (top panel; the residue index is nonlinearly scaled for better visualization) with the more detailed definitions from (Shi W. et al., 2023, *Nature* 619, 403-409) (bottom panel; coloring of the main segment preserved from the top panel). (B) Predicted frustration calculated from frustratometer server (http://frustratometer.qb.fcen.uba.ar/) for the three chains of the S2 subunit. Density of highly frustrated (or destabilizing) contacts for each residue, calculated using a 0.5 nm cutoff radius, for the prefusion (red) and postfusion (black) states, and plotted vs residues as numbered in the full S protein. The domains are marked above the plots. Overall, the postfusion state shows lower frustration than the prefusion state, suggesting it is more stable. We use this prediction to justify using a Cα-SBM of the postfusion state in our simulations.

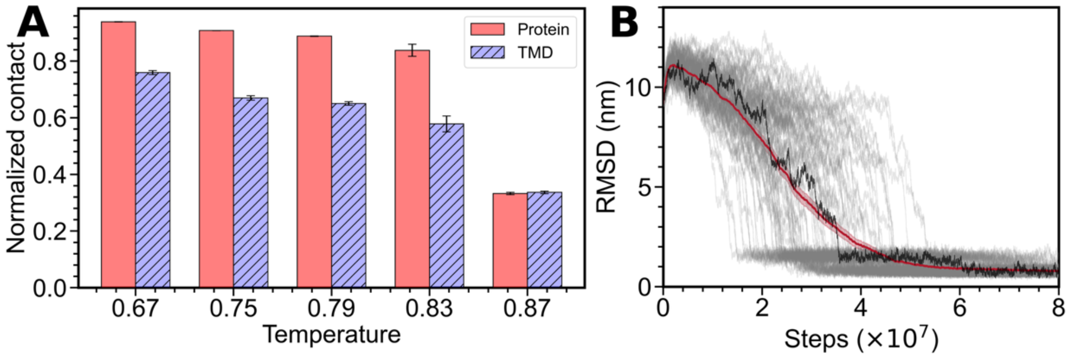


**Figure S3: Choosing a simulation temperature.** (A) Normalized fraction of native (postfusion structure) contacts as a function of temperature for the protein (S2, 2410 contacts) and the TMD residues (protein contacts involving TMD residues; 99 contacts). As the reduced temperature increases, there is a higher contact loss. (B) Using (A) a simulation (reduced) temperature of 0.75 was chosen, and 100 trajectories were simulated (see **Figure 2**). Root mean square deviation (RMSD) with respect to the postfusion state as a function of time is shown for individual trajectories (gray) with one trajectory highlighted (black). The mean of the data (red line) is also shown, with the error of the mean marked by the shaded region around the mean.

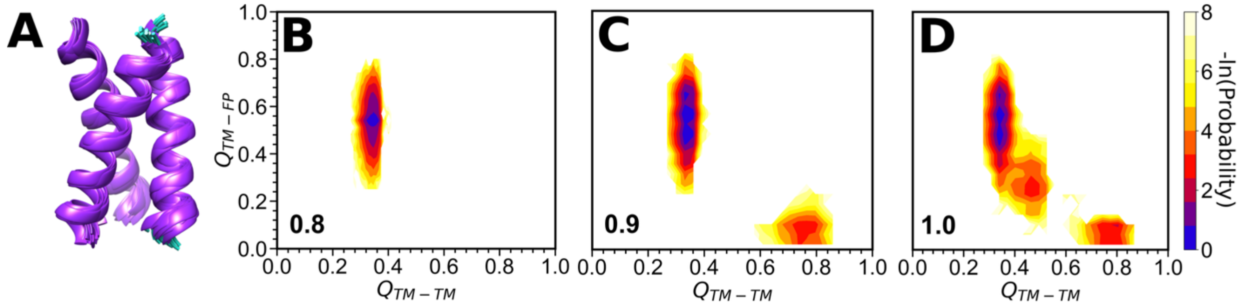


**Figure S4: Choosing the strength of NMR derived contacts for the TMD.** (A) The NMR ensemble of the isolated TMD (PDB ID: 7LC8) (B-D) 2D-free energy surfaces with fraction of TMD-FP contacts (38 total; y-axis) and fraction of TMD-TMD contacts derived from the NMR structure (111 total; x-axis). For these calculations, snapshots were extracted from post-conversion converged parts of conformational transition trajectories. Simulations were performed with NMR contact strengths of (B) 0.8ε (where ε is the strength of other contacts), (C) 0.9ε, and (D) 1.0ε. Since TMD-TMD contact strengths of up to 0.8ε do not perturb the final postfusion state, this strength was used for the simulations.

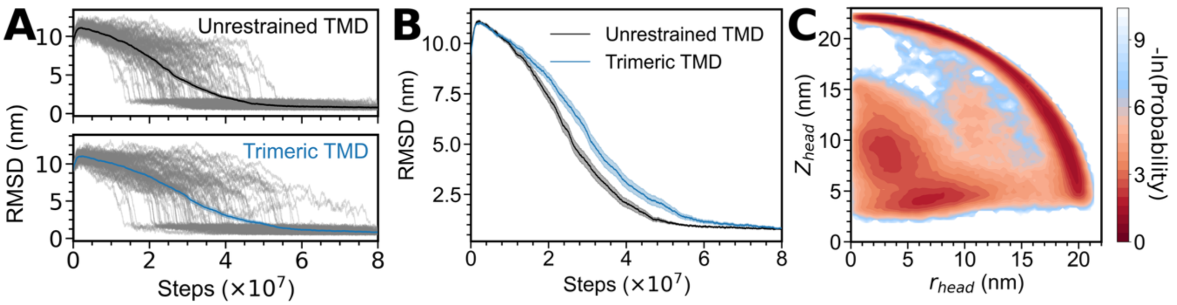


**Figure S5: Additional comparison of the Unrestrained TMD and Trimeric TMD models.** (A) (A) RMSD from the unrestrained TMD (upper panel) and trimeric TMD (lower panel) models. Individual simulation data (100 replicates; gray lines) plotted as a function of time. The mean is shown. (A-B) Black is used to depict the unrestrained TMD, and blue to depict the trimeric TMD models. (B) A comparison of the mean RMSD from (A). The shaded regions represent the error of mean estimation. The trimeric TMD model converts slower. (C) Two-dimensional free-energy surface with $Z_{head}$ and $r_{head}$ as coordinates. See **Figure 2** for details of the coordinates and **Figure 2D** for comparison with the Unrestrained TMD model. Specifically, the addition of the TMD contacts stabilizes a state near ($r_{head}$ ~ 3 nm, $Z_{head}$ ~ 8 nm).

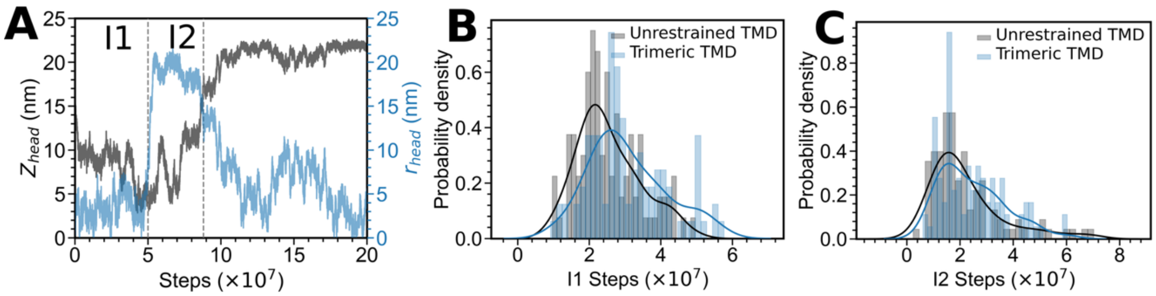


**Figure S6: A comparison of the sampling of intermediates I1 and I2 in the Unrestrained TMD and Trimeric TMD models.** (A) $Z_{head}$ (grey) and $r_{head}$ (blue), calculated from a representative trajectory and plotted as a function of time. The protein is in I1 if $Z_{head}<15$ nm and $r_{head}<10$ nm, and in I2 if $Z_{head}<15$ nm and $r_{head}>10$ nm. (B, C) Probability density distributions of the number of steps sampled in the intermediate states (one data point per trajectory) in the unrestrained TMD (grey) and the trimeric TMD (blue) models. (B) I1 (C) I2. Both I1 and I2 are on an average sampled longer in the trimeric TMD model.

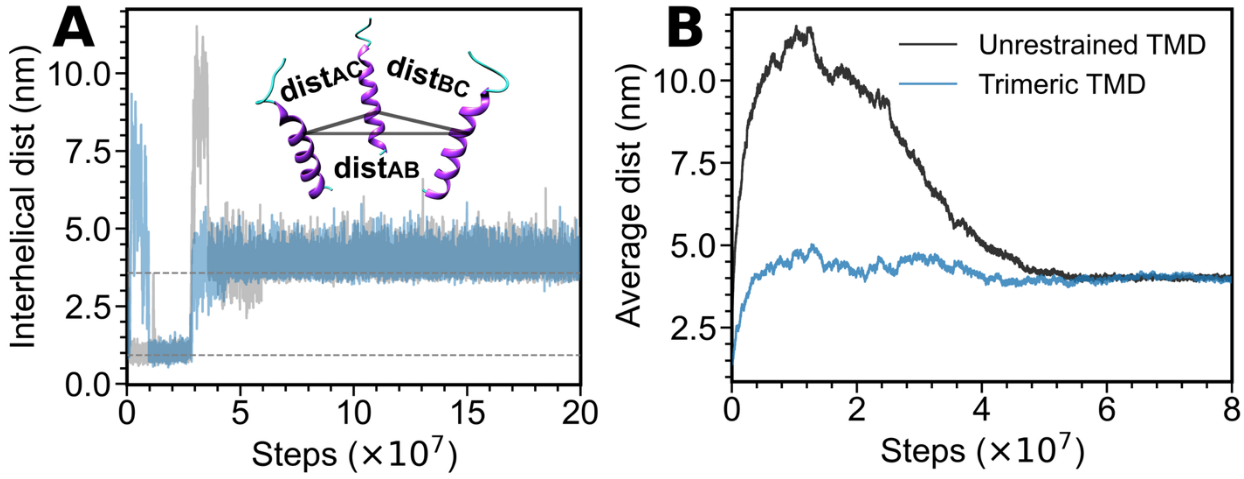


**Figure S7: Definition of TMD dissociation and a comparison between the Unrestrained TMD and Trimeric TMD models.** (A) Representative inter-helical distance traces from two individual trajectories. The inset illustrates how the average inter-helical distance is defined: the distance between the centers of mass of any two helices X and Y in the TMD, $dist_{XY}$, is calculated, and the average of the three marked distances is the average inter-helical distance. Gray dashed horizontal lines mark the prefusion (0.93 nm) and postfusion (3.5 nm) values of the average interhelical distance. A TMD is considered to be dissociated or "open" when the average interhelical distance exceeds the prefusion state value and remains above it. (B) The average interhelical distance from (A) averaged over the 100 trajectories of the unrestrained TMD (black) and the trimeric TMD (blue) models shows much smaller average interhelical distances in the trimeric TMD model.

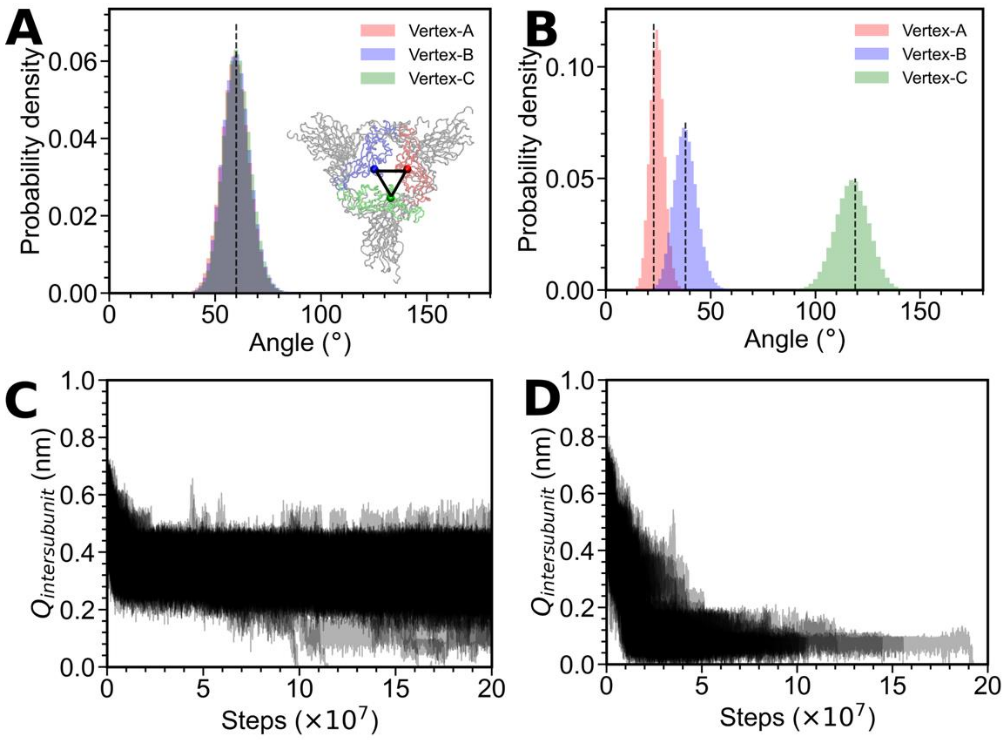


**Figure S8: Analyses of the S1-closed RBD and S1-open RBD models.** (A, B) The same set of 100 spike (S1 and S2) structures with all closed RBDs was used to initiate the S1-closed and S1-open RBD simulations. The amino acids numbered 445 from each of the three (A, B, C) RBDs form a triangle (A, inset). The angle at each of the vertices of this triangle was calculated from all simulation snapshots after discarding initial 10000 steps. (A) S1-closed RBD model: The three RBDs remain closed and symmetric, and the probability density of the three angles across the simulation trajectory overlaps and merges into the grey distribution centered at 60° (black dashed line; native state). (B) S1-open RBD model. One RBD opens, and the vertex angles diverge into three distributions (red, blue, green). The peaks of the distributions closely match the native values (22°, 38°, and 119°) calculated from the S1-open RBD structure. This implies an accurate encoding of the structure in the SBM. (C, D) The fraction of inter-subunit (between S1 and S2) contacts ($Q_{intersubunit}$) as a function of time, plotted for all 100 trajectories. (C) S1-closed RBD model (355 native S1-S2 contacts). Many inter-subunit contacts are retained, and S1 is completely shed ($Q_{intersubunit}$=0) within the simulation time in only 5 out of 100 trajectories (D) S1-open RBD model (291 native S1-S2 contacts). $Q_{intersubunit}$ becomes 0, and S1 is shed within the simulation time in all trajectories.

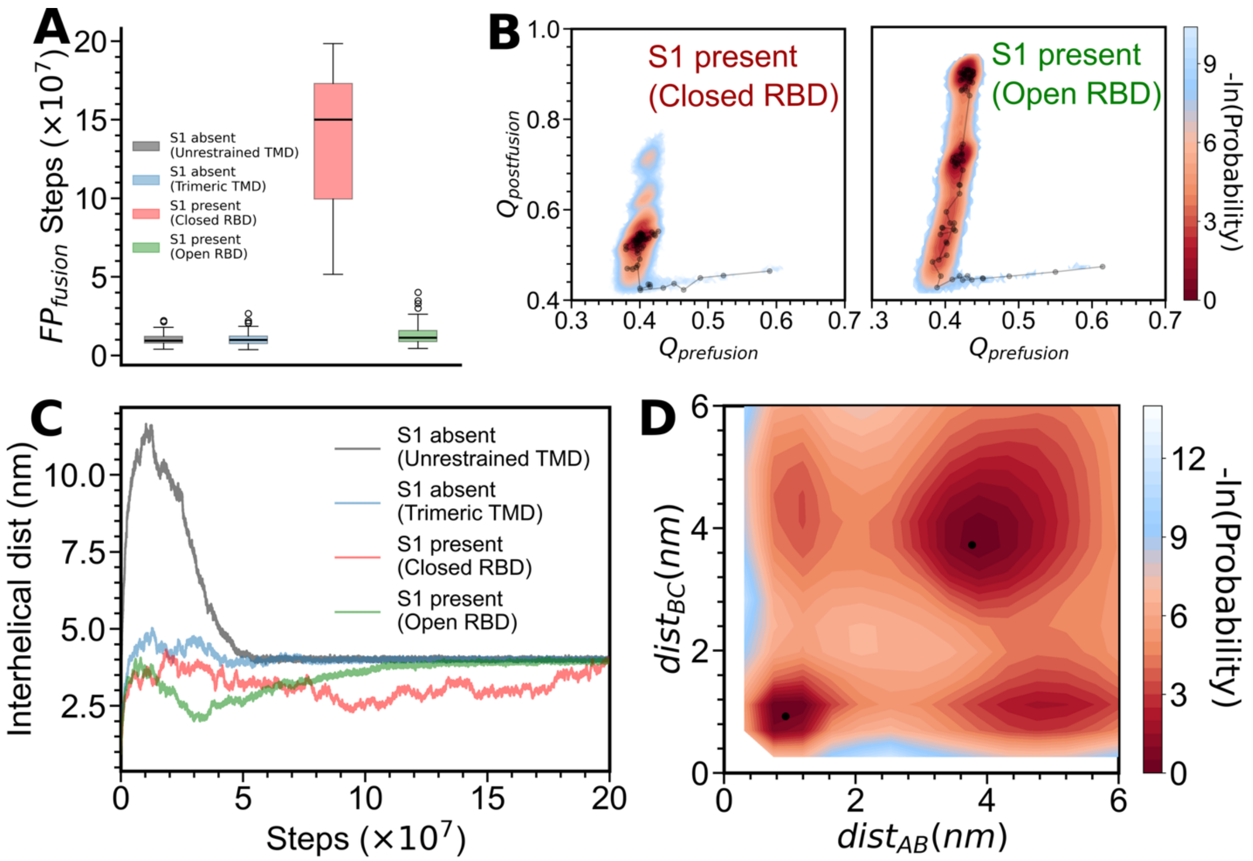


**Figure S9: Additional analysis of S1 containing simulations.** (A) $FP_{fusion}$ is the number of steps needed for the fusion head to form, or the average distance between every pair of centers of mass of the three fusion peptides, $d_{FP} < 1.27$ nm. Since S1 sheds in very few simulations, $FP_{fusion}$ (S1-closed RBD) >> $FP_{fusion}$ (other models). (B) Two-dimensional free energy surfaces with the fraction of formed prefusion contacts (2344 native contacts) and the fraction of formed postfusion contacts (2410 native contacts) as the two coordinates plotted using conformations of the S2 subunit obtained from 100 trajectories using the S1-closed RBD (left panel) and S1-open RBD (right panel) models. A representative trajectory is overlaid upon the plots and shows that not all S1-closed RBD transitions may be complete. (C) Average interhelical distance between the TM helices (see **Figure S7** for the definition) plotted as a function of time for the different models used. (D) Two-dimensional free energy surface for the S1-open RBD model with the interhelical distances $dist_{BC}$ and $dist_{AB}$ as the two coordinates (see **Figure S7A** for the definition). Data from all 100 complete trajectories were used for this calculation, unlike in **Figure 4D**, where only the portion of the trajectory with S1 present was considered.

**DATA S1:** S2-prefusion PDB

**DATA S2:** S2-postfusion PDB

**DATA S3:** Contact list for the S2-prefusion SBM. Atom numbers 1-1248 correspond to the three chains spanning residues 834-1249.

**DATA S4:** Contact list for the unrestrained TMD model. Atom numbers 1-1248 correspond to the three chains spanning residues 834-1249.

**DATA S5:** Contact list of prefusion trimer specific NMR contacts. The atom numbering corresponds to the same sequence as in DATA S4.

**DATA S6:** PDB structure of the S1/S2 complex with all RBDs in the closed state.

**DATA S7:** PDB structure of the S1/S2 complex with one RBD in the open state.

**DATA S8:** Contact list for the S1/S2 complex with all closed RBD prefusion structure. Atom numbers are sequentially assigned from 1 to 3303, corresponding to residues 1-685 and 834-1249.

**DATA S9:** Contact list for the S1-present closed RBD model. Atom numbers are sequentially assigned from 1 to 3303, corresponding to residues 1-685 and 834-1249.

**DATA S10:** Contact list for the S1-present open RBD model. Atom numbers are sequentially assigned from 1 to 3303, corresponding to residues 1-685 and 834-1249.

**MOVIE S1:** S2 conformational transition captured using a trimeric TMD model. The implicit viral membrane and domains are highlighted. The host membrane is shown for visualization only and is not included in the simulations.
